# Actin and microtubules position stress granules

**DOI:** 10.1101/2023.06.04.543599

**Authors:** Thomas J. Böddeker, Anja Rusch, Keno Leeners, Michael P. Murrell, Eric R. Dufresne

## Abstract

Membraneless organelles, composed of protein and nucleic acids, alter the biochemical and physical landscape of the cell. While specific membraneless organelles are found in stereotypical locations, little is known about the physical mechanisms that guide their positioning. Here, we investigate how stress granules, a type of cytoplasmic membraneless organelle, establish their stereotypical perinuclear positioning. We find that actin and microtubules play complementary roles. Lamellar actin confines stress granules, and its retrograde flow drives them toward the cell center. Microtubules, in turn, adhere to stress granules through capillary interactions, which tend to concentrate stress granules in micro-tubule rich regions near the nucleus. Similar physical mechanisms are likely to play a role in the positioning of other membraneless organelles.

## I. INTRODUCTION

To survive and proliferate, cells need to regulate a vast number of biochemical reactions. To facilitate this daunting task, cells compartmentalize many of these reactions within organelles, including classical membrane-bound organelles (such as mitochondria or endoplasmic reticula), or emerging membraneless organelles (such as the nucleolus in the nucleus or stress granules and p-bodies in the cytosol) [1]. The function of membrane-bound organelles has been shown to depend on their position within the cell, with impact on signaling, polarization, or cell growth [2]. Their positioning can be assisted by active mechanisms, such as motor-driven transport along microtubules [2–5].

Less is known about motion and positioning of membraneless organelles in the cytosol. These biomolecular condensates are typically composed of protein and mRNA [1, 6]. Stress granules (SGs) are cytoplasmic membraneless organelles that serve as dynamic sites of mRNA sorting. They are important for reorganization of translation under biological stress (*e.g.* exposure to toxic chemicals or heat) [7–9]. Because the formation of SGs can easily be induced, they are a convenient model system to study the appearence and localization of membraneless organelles in the cytosol. A key protein for SG formation under oxidative stress, such as caused by exposure to arsenite, is G3BP1 [10, 11]. In unperturbed cells, G3BP1, as well as other SG components, are dispersed throughout the cytosol. Upon exposure to stress, these molecules condense to form SGs throughout the cytosol before migrating toward the cell center [7, 12–14]. Ultimately, SGs are found in the microtubule-rich perinuclear region of the cytoplasm [15, 16]. As SGs travel toward the nucleus, they coalesce and grow, resulting in several micron-sized granules within tens of minutes of exposure to stress [13, 14, 17].

Cytoskeleton components are obvious suspects for the active control of the position of membraneless organelles. Microtubules have been suggested to aid stress granule formation by acting as tracks for active transport of granule components through motor proteins [18–22] and by encouraging droplet fusion through mixing of the cytosol during their dynamic instability [23]. Early studies reported that microtubules were neccessary for the formation of stress granules [18, 19], while more recent ones found that SGs simply have a reduced size in the absence of a microtubule network [13, 14, 24]. A microtubule-associated motor, dynein, was reported to have no impact on SG formation [14]. On the other hand, p-bodies could play a role, as they are known to both associate with SGs [10, 25] and to be actively transported along microtubules [26]. Overall, these studies suggest that microtubules and associated proteins could influence SG positioning. On the other hand, disruption of the actin network has been reported to have no effect on SG formation [19, 22].

Here, we investigate the potential impact of actin and microtubules on the positioning of SGs. Combining quantitative structural measures in fixed-cells, with live-cell dynamics and cytoskeletal perturbations, we identify distinct roles for actin and microtubules. Lamellar actin confines stress granules, and its retrograde flow transports them toward the cell center. Once there, microtubules guide SGs to their final location with attractive capillary interactions. These generic physical mechanisms are likely to play a role in the positioning of other membraneless organelles.

## II. RESULTS

### A. Birth and maturation of stress granules

To observe the dynamics of SGs throughout the cell’s stress response, we p imaged live human epithelial U2OS cells on a confocal microscope. Figure 1 shows representative fluorescence images of a U2OS RDG3 cell, expressing GFP-tagged G3BP1 [27], exposed to 150 µM arsenite at time zero. Let us first compare the cell before and after arsenite treatment. Figure 1 **a** shows the distribution of G3BP1 at the onset of arsenite treatment and after 90 min. In accordance with previous results, we find that G3BP1 is distributed throughout the cytosol at the onset of stress but localizes in micron-sized SGs in the perinuclear region after the stress response of the cell. Formation of SGs is not the only change in the cell upon exposure to stressful conditions – the morphology of the actin network also changes. We visualize filamentous (f)-actin using SPY650-FastAct (Spyrochrome) and find that, while f-actin is initially most pronounced along the cell periphery, the actin network has significantly contracted after 90 min of arsenite treatment, see Fig. 1 **b**, hinting at a role for actin in directing SG motion. The full time series of the experiment is shown in Fig. 1 **c**. We find that SGs nucleate out of a pool of G3BP1 after about 7.5 min of arsenite treatment^1^. SGs appear to rapidly move towards the cell center within a few minutes after formation. Throughout this motion, SGs coalesce leading to fewer, larger, granules closer to the cell center after about 30 min after exposure to arsenite. At longer times, SGs continue to move inward and fuse, albeit at an apparently slower pace, leading to little change in SG distribution and size after about 60 min. Note that small SGs remain under and above the cell nucleus.

**FIG. 1.**
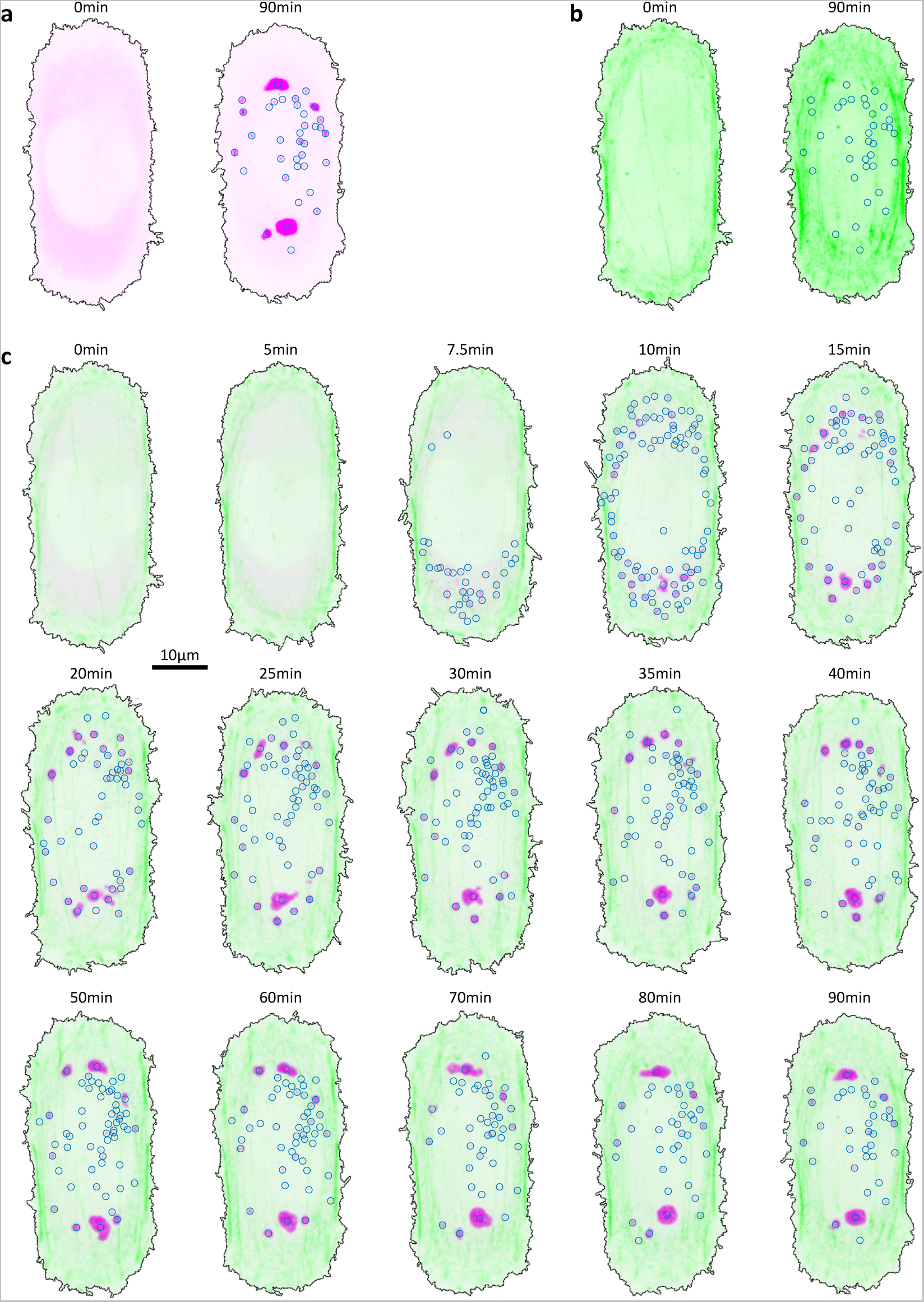
Stress granule dynamics in an exemplary experiment with a U2OS RDG3 cell exposed to 150 µM arsenite at 0 min. Images show the maximum projection of a confocal stack. Color saturation is consistent within cells of a given panel. G3BP1 is shown in magenta, f-actin in green. **a**, comparison of G3BP1 at the onset of arsenite treatment and after 90 min. The blue circles indicate the position of detected SGs. The black line indicates the cell outline. **b**, comparison of actin at the onset of arsenite treatment and after 90 min. **c**, time series of an exemplary experiment. First SGs appear after 7.5 min of arsenite treatment. These images or single timepoints from Supplementary Movie 1.

SGs coalesce and migrate over time, showing initially fast dynamics that slow down over time, as qualitatively observed in the single cell. In order to quantify these observations, we record the position and volume of granules within the confocal stack of the G3BP1 channel over time. The circles in Fig. 1, indicate the centroid of registered SGs. Figure 2 a - c show the number of SGs, N*_SG_*, the mean granule volume, ⟨V ⟩, and the total granule volume, V_total_, for the cell shown in Fig. 1. For the time axis, we introduce t*_SG_*, which indicates the time after the first SGs are detected in the cell. For this cell, t*_SG_* = 0 was 7.5 min after onset of arsenite treatment.

**FIG. 2.**
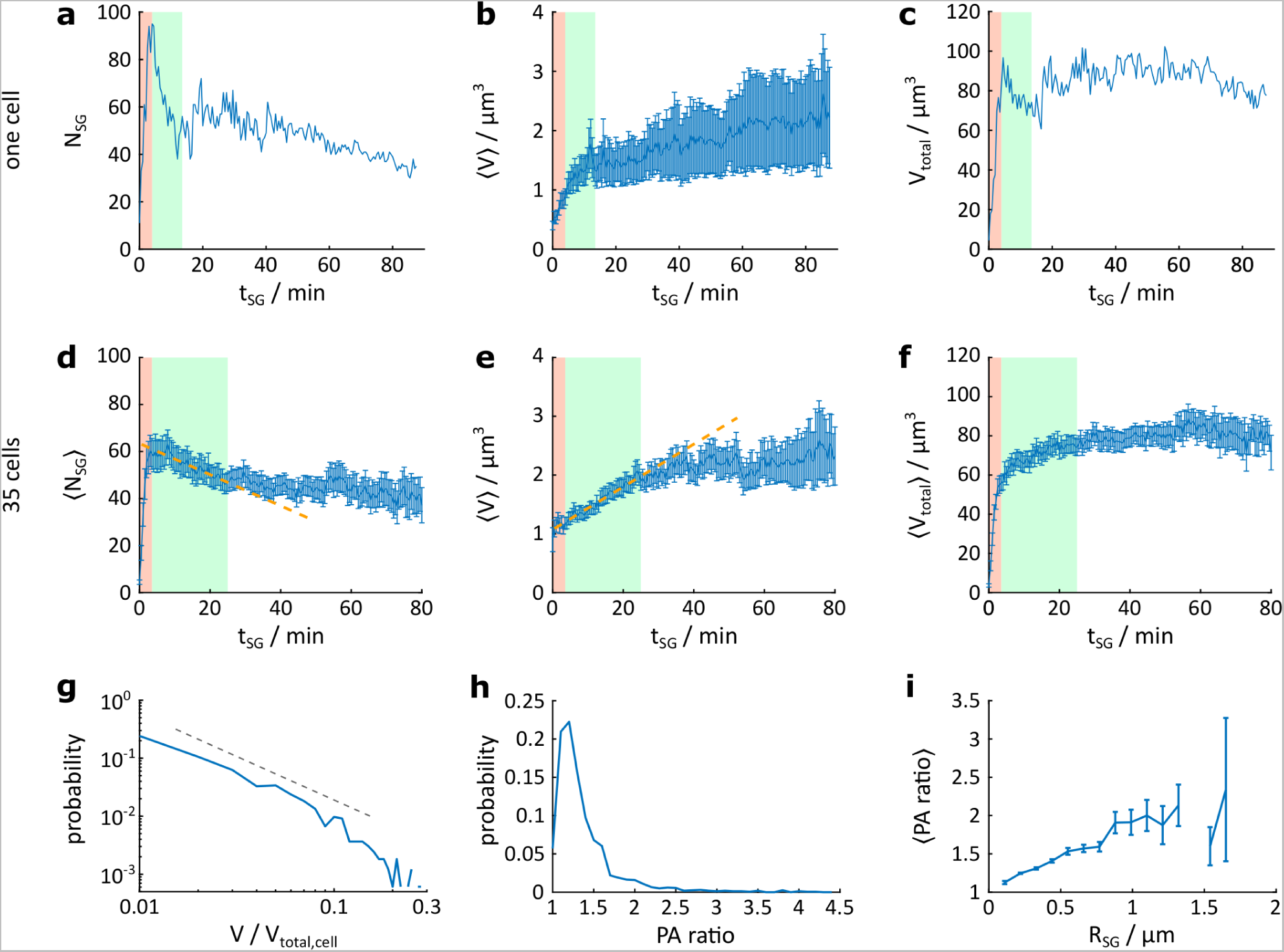
Statistics of SG formation over of time after detection of the first granule at *t_SG_* = 0 min. **a**, the number *N_SG_* of detected SGs in the cell shown in Fig. 1. **b**, the average volume ⟨*V* ⟩ of SGs in this cell, the error is given as the standard error. **c**, the total volume *V*_total_ of SGs in this cell. The nucleation and growth phase is shaded in red and the coalescence phase is shaded in green. **d**, ⟨*N_SG_*⟩ averaged over *N* = 35 cells, the error is given as the standard error. The orange dashed line is a guide to the eye. **e**, ⟨*V* ⟩ averaged over *N* = 35 cells, the error is given as the standard error. **f**, ⟨*V*_total_⟩ averaged over *N* = 35 cells, the error is given as the standard error. The red shaded area indicates the phase separation regime, the green shaded area the fast-ripening regime. **g**, probability distribution of the SG volume *V* at *t_SG_* = 25 min with 1645 SGs considered. *V* is normalized by *V*_total_ for each cell. The dashed lines indicates a slope of -1.5. **h**, probability distribution of the principal axis (PA) ratio for granules at *t_SG_* = 25 min. **i**, mean PA ratio, as a function of *R_SG_* at the same time point. Error bars indicate the standard error.

This quantification highlights three phases of SG maturation, see Fig. 2. The first phase, which we call nucleation and growth (labelled in red), lasts for about 5 min after the first SGs appear and is characterized by a rapid increase in N*_SG_*, ⟨V ⟩, and V_total_. In the second phase, which we call coalescence (labelled in green), V_total_ is stable, indicating that further changes in number and volume of SGs are dominated by ripening processes, not by further recruitment of molecules to SGs. Indeed, we find a rapid decrease of N*_SG_* that coincides with a quick increase in ⟨V ⟩, indicative of coalescence. After about t*_SG_* = 15 min, SGs enter a slow-ripening phase, where N*_SG_* and ⟨V ⟩ only change marginally: N*_SG_* slowly decreases and ⟨V ⟩ slowly increases.

To more rigorously quantify this stochastic process, we combine experiments across N = 35 cells. To enable meaningful data-pooling, all cells are plated on fibronectin-coated patterns in the shape of a rectangle with two semi-circular caps on a coverslip [28]. Spatially aligning micrographs of multiple cells and averaging the intensity of a given channel, we can construct time-resolved ensemble averages, see Supplementary Fig. 14 and Supplemental Movie 2. These show that the overall spatio-temporal dynamics of SG birth and maturation are highly reproducible. Next, we detect SGs in each cell separately, and pool the results for all 35 cells in Fig. 2 **d - f**. The previously described stages (nucleation and growth, coalescence, and slow-ripening) remain clear in the ensemble-averaged data, but with slightly different timing. On average, the nucleation and growth regime persists for about 5 min after formation of first SGs. This is followed by a phase of coalescence that lasts until about t*_SG_* = 25 min. Afterward, SGs ripen slowly, decreasing in number and increasing in size. Ultimately, stress granules occupy about 6% of the cytosol^2^.

The distributions of SG size and shape provide further insights into their growth and maturation. Figure 2 **g** shows the probability distribution of the granule volume V for 1645 granules at t*_SG_* = 25 min. We find that the volume distribution is consistent with a truncated power-law distribution with an exponent of about -1.5, as reported for the volume distribution of condensates in the cell nucleus [29, 30]. SGs are typically not spherical. To quantify this, we used the principal axis (PA) ratio, *i.e.* the ratio of the lengths of the major and minor principal axes of an ellipse fitted to each granule. The distribution of PA ratio at t*_SG_* = 25 min, shown in Fig. 2 **h**, reveals that only about 6% of all SGs are indistinguishable from spheres. Most granules are slightly deformed out of a spherical shape and have a PA ratio between 1 and 1.5. The remaining 19% of SGs have a PA ratio larger than 1.5, reaching values beyond 4. Further, the mean PA ratio increases with SG size, see Fig. 2 **i**. These deformations can relax over time and are thus not indicative of changes in SG material properties due to hardening (see Supplementary Movie 1). Since SGs are deformed from the spherical shape favored by surface tension, additional forces must be acting upon them.

### B. Interactions of Stress Granules with Microtubules and Actin

The cytoskeleton is a complex network spanning the cytosol composed of actin, microtubules, and various intermediate filaments. The cytoskeleton has heterogeneous material properties [31] – its organization and mesh-size vary throughout the cytosol [32, 33].

SGs interact with microtubules, as found in previous studies. While early studies suggested that the observed co-localization of SGs focused on motor proteins [18–22], recent work has highlighted the importance of wetting interactions between microtubules and SGs that lead to a significant enhancement of microtubule density around SGs. These wetting interactions are a consequence of SGs’ native surface tension. Tubulin dimers, the molecular building blocks of microtubules, have no preference for either SG or cytosol. Therefore, they can act as weak Pickering agents, adsorbing to the SG interface. While the adhesion strength of isolated dimers is less than k*_b_* T, polymerized microtubules can bind strongly, with an adhesion strengths up to 50 k*_b_*T/ µm [24].

Building upon these findings, we compare the interactions of SGs with microtubules and f-actin. We take a structural approach, quantifying correlations of SGs with actin and microtubules, using confocal stacks of U2OS cells fixed after 90 min of exposure to arsenite. All these cells are plated on the same patterned substrate and can therefore be spatially registered. Averaging over about 300 cells, we construct *reference cells* that capture the typical distribution of f-actin, *β*−tubulin and G3BP1 in these cells, see **Fig. 3 a - c**. Note that reference cells are resolved in three dimensions, see Supplementary Fig. 9. We find that the actin and microtubule networks are dense in different regions of the cell, with microtubules predominantly in the perinuclear region of the cell and actin on the periphery. High intensity of G3BP1 tends to co-localize with high *β*−tubulin intensity, while actin-rich regions show very low G3BP1 signal. For detailed information on construction of the reference cells refer to Materials and Methods and [24]. An example of the actin, SG, and microtubule channels for a single cell is shown in Supplementary Fig. 8 **a** and **b**.

**FIG. 3.**
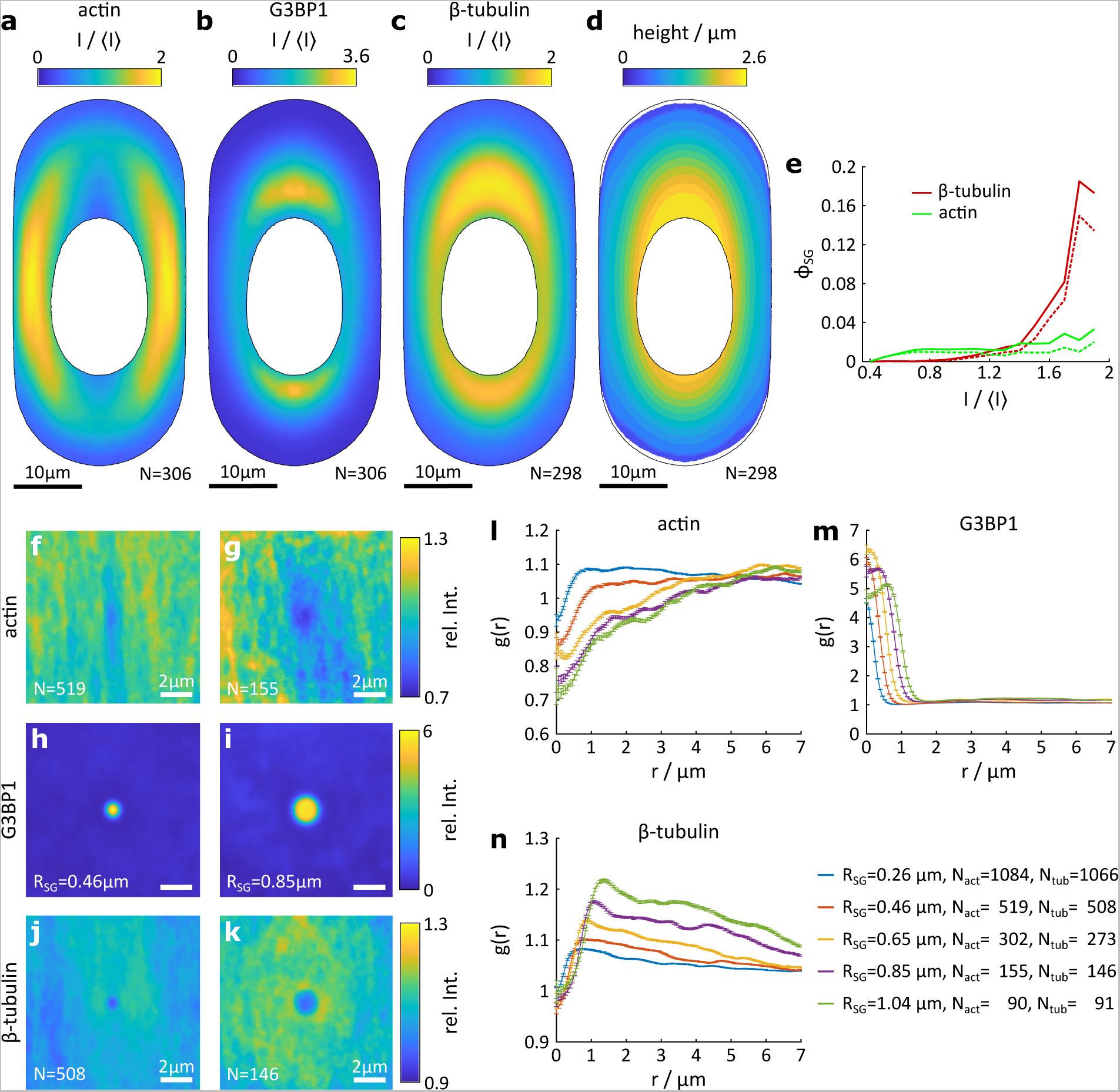
Stress granules are attracted by microtubules and repelled by actin. **a**, **b**, **c**, reference cell of the actin, G3BP1 and microtubule channels averaged over *N* cells fixed after 90 min of exposure to arsenite. The intensity is normalized by the average intensity in each individual cell and channel ⟨*I*⟩. The black line indicates the extent of the cytosol. **d**, height profile of the reference cell. **e**, volume fraction *ϕ_SG_* of stress granules in regions of the cell with a given intensity of the actin and *β*−tubulin channel. Solid lines indicate *ϕ_SG_* calculated using the number of registered voxels of SGs, dashed lines show *ϕ_SG_* calculated based on the radius of each granule given by the location of the maximum gradient in the G3BP1 channel. **f**, **g**, distribution map of actin around stress granules of roughly spherical shape (principal axis ratio below 1.5) and with a radius in the interval from 0.39 to 0.52 µm (indicated as *R_SG_* = 0.46 µm) and from 0.78 0.91 µm (indicated as *R_SG_* = 0.85 µm). *N* gives the number of contributing SGs. **h**, **i**, distribution map of G3BP1 for small and larger SGs. **j**, **k**, distribution map of *β*−tubulin for small and larger SGs. **l**, radial distribution function *g*(*r*) of actin for granules of varying size. **m**, **n**, *g*(*r*) for G3BP1 and *β*−tubulin respectively. The control for the data in panel **g** - **o** is shown in F^9^ig. 10 in the Supplement.

Actin-rich regions in the cell periphery are thinner than the region around the cell nucleus, where most microtubules are located. The apparent affinity of SGs towards microtubules and avoidance of actin might therefore be impacted by volume effects. To probe for such effects, we quantify the cell height through a low intensity threshold of the *β*−tubulin channel as shown in Fig. 3 **d.** We find that the cell height continuously increases from the cell periphery towards the cell nucleus. The maximum of G3BP1 intensity, however, coincides with the highest intensity in the microtubule network and not with the regions of largest cell height. Further, side views of the reference cells are shown in Supplementary Fig. 9 reveal that the alignment of G3BP1 with *β*−tubulin also holds across the thickness of the cell. These observations rule out a dominant role for cell thickness as the main driving force for the positioning of SGs.

To probe the affinity of G3BP1 to actin and *β*−tubulin, we next explore the dependence of the local SG volume fraction, *ϕ_SG_*, on the local intensity of f-actin and *β*−tubulin, I/⟨I⟩, see Fig. 3 **e**. At very low I/⟨I⟩ of actin, that is at the very cell edge, no SGs are present. Regions of intermediate actin intensity have a roughly constant *ϕ_SG_* of about 0.02, which only increases slightly to about 0.03 in regions of high actin intensity. For *β*−tubulin, we find a very different behavior. *ϕ_SG_* increases with increasing tubulin intensity, reaching surprisingly high values of about 0.16 in the regions of highest *β*−tubulin intensity, suggesting a strong affinity of SGs for microtubule-rich regions. A similar observation is made when correlating the local intensity of G3BP1 with actin and *β*−tubulin intensity. The results are shown in Supplemental Fig. 8 **c** and **d**. We conclude that SGs have a strong tendency to co-localize with *β*−tubulin-rich regions of the cell and tend to avoid the actin-rich cell periphery.

While we have broadly quantified the tendency for SGs to inhabit the same parts of the cell as actin and microtubules, the above analyses do not resolve the interactions of individual SGs with cytoskeletal filaments. To that end, we quantify local structural correlations of the microtubule and actin network around SGs. This correlation analysis leads to *distribution maps* shown in Fig. 3 **f** - **k**. Distribution maps show the average intensity of a given channel around SGs of selected size and shape, here SGs with a roughly spherical shape (having a PA ratio below 1.5) are binned by their radius in steps of 0.195 µm. Throughout the construction, individual images contributing to distribution maps are normalized against the respective reference cell, see Materials and Methods and [24]. Distribution maps consequently highlight average changes of the network around individual stress granules. A value of one means the observed intensity is identical to the reference cell, while values above (below) one reveal a scaled increase (decrease) of the intensity. A computational negative control for this analysis is shown in Supplemental Fig. 10.

The distribution maps for G3BP1 (Fig. 3 **i** and **j**) show spherical regions of elevated G3BP1 intensity with a radius corresponding to the radius of the contributing SGs. To quantitatively compare SGs of different size, we use the *radial distribution function* g(r), the azimuthal average of the distribution map. g(r) for G3BP1 is shown in Fig. 3 **n** and shows the expected outward shift as the R*_SG_* increases.

SGs show different affinities towards actin and microtubules. To see this, we compare the network structure and intensity of actin and microtubules around SGs. Our correlation analysis reveals that actin retains its normal configuration, *i.e.* a stripe like morphology originating from stress fibers, around small SGs (Fig. 3 **g**), while SGs seem to expel or avoid actin fibers, with relative intensities below one at the position of the granules. Further, small SGs are found in regions of higher than expected actin density, with up to 1.1 times higher actin intensity around the smallest SGs. Larger SGs are found in regions of lower than expected actin density, with the largest granules present in regions of only 0.7 times the expected actin density (Fig. 3 **h**). Larger SGs further have a zone of depleted actin intensity around them that extends over several granule radii. g(r) for actin shown in Fig. 3 **m** clearly shows the emergence of an actin exclusion zone for SGs of increasing radius. These data suggest the the presence of a dense actin network might limit granule size.

For the microtubule channel, we find an enhancement in *β*−tubulin intensity around SGs of all sizes. This enhancement is centered on SGs and increases in magnitude as SG size increases (Fig. 3 **k** and **l**). Further, it decays roughly linearly and extends over several granule radii away from the SG. The relative *β*−tubulin intensity within SGs, however, is close to one for all SGs. The enhancement of microtubules increases with increasing SG size, as visible in the g(r) (Fig. 3 **o**). We previously interpreted these observations with adhesive forces between SGs and microtubules [24]. Interestingly, the extent of the enhancement in *β*−tubulin intensity around larger SGs roughly corresponds to the depletion zone in the g(r) for actin. This suggests possible steric repulsions between the two networks.

Overall, these results suggest a short-ranged repulsion between SGs and actin, and adhesion of SGs and microtubules. While this static correlation analysis is informative of interactions between SGs and cytoskeletal filaments, we need to return to live-cell experiments to visualize how these interactions could ultimately determine the position of stress granules.

### C. Time-resolved Transport of Stress Granules

While SGs form throughout the cytosol, they are strongly localized to the perinuclear region after about t*_SG_* = 25 min. This shift in SG distribution toward the perinuclear region occurs primarily during their nucleation and growth and coalescence phases, see Fig. 1 and Supplementary Fig. 14. The inward transport of SGs is reminiscent of the suggestive flow of actin in the lamellopodium [33, 34].

In order to investigate a potential coupling of SG-transport with motion of the actin network, we performed particle image velocimetry (PIV) of actin and G3BP1. To quantify the average flow speed and direction of both f-actin and G3BP1, we construct *PIV ensembles*, by averaging PIV results from individual cells over time bins of 2.5 min and, again, averaging these results over 35 cells. For details on the PIV analysis, refer to Materials and Methods. Figure 4 **a** and **b** show the PIV ensembles for the actin and G3BP1 channels at t*_SG_* = 10 min. The full time series of the PIV ensemble is shown in Supplementary Movie 3. We find that actin flow is directed towards the cell center and is especially pronounced in the lamellar regions at the curved ends of the cell. The flow of G3BP1 is also, on average, inward but exhibits larger fluctuations.

**FIG. 4.**
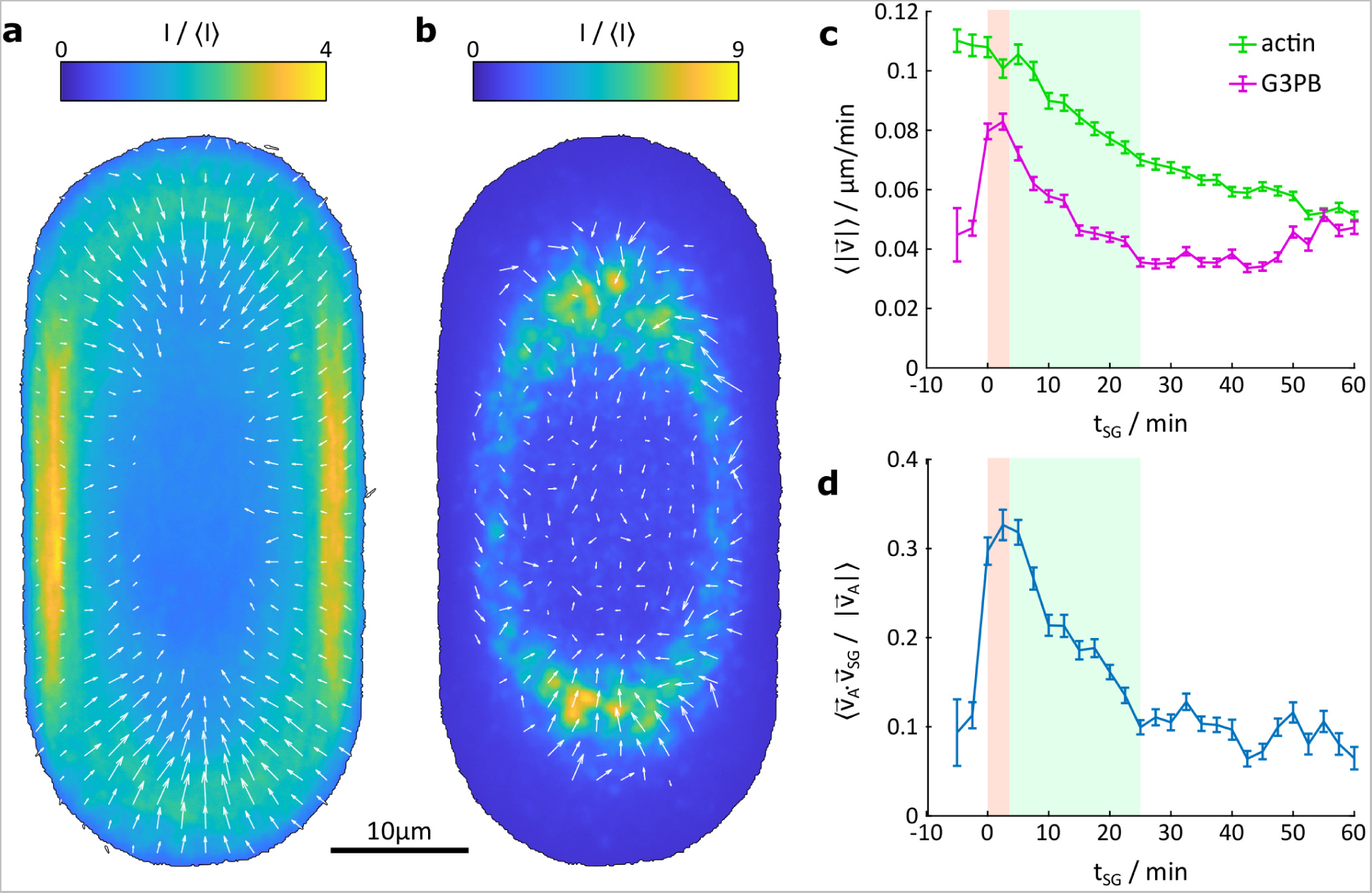
Actin and G3BP1 flow are aligned. **a**, PIV ensemble for *t_SG_* = 10 min for the actin channel overlain with the actin ensemble averaged over the same data. **b**, the corresponding PIV ensemble for G3BP1. **c**, the mean velocity of actin and G3BP1 throughout the cell over time. Error bars show standard error. The phase separation and fast-ripening regime are shaded in red and green (compare Fig. 2). **d**, the alignment calculated as the dot product between actin and G3BP1 flow over time normalized by the amplitude of the actin flow. Error bars show standard error.

When stress granules first form, they flow with the actin network. The cell-wide average speeds, |⃗v|, of actin and G3BP1 are shown in Figure 4 **c**. The average actin speed is initially steady at 0.11 µm/min. At the same time, the maximum speed is higher, reaching 0.29 µm/min, see Supplementary Fig. 11 **a**. These values are consistent with previous reports of actin flow rates, [34, 35]. When stress granules first appear, their speed nearly matches that of actin, with the cell-wide average the of G3BP1 channel reaching 0.08 µm/min, and a maximum speed of 0.25 µm/min (Supplementary Fig. 11 **b**). The flow of actin and G3BP1 are well aligned, as shown in Fig. 4 **d**. As stress granules start to coalesce, the average speeds of the actin and G3BP1 channels slow down (Fig. 4 **c**), and become less well-aligned (Fig. 4 **d**). By the end of the coalescence phase (around t*_SG_* = 25 min), there is no significant flux of G3BP1.

The observed actin flow has two contributions, lamellar retrograde flow [34, 35] and a stress-induced net contraction of the actin cytoskeleton [36]. The latter is most clearly visible by comparing the actin reference cells across time, see Supplementary Fig. 14. We find a contraction speed of about 0.1 µm/min along the mid-line of the cell, see Supplementary Fig. 12. Contraction occurs at this constant speed from t*_SG_* = 0 to 40 min, then fully stops after about t*_SG_* = 50 min, during the slow-ripening phase.

Overall, we find that motion of SGs and actin flow are positively correlated throughout the ripening process with highest speeds and alignment throughout early stages of the cell’s stress response when SGs are still present in the lamellar region of the cell. Our data suggest that retrograde flow of actin leads to rapid transport of SGs out of the lamellar region of the cell and that subsequent contraction of the actin network induces further inward displacement of SGs.

### D. Emerging Model and Perturbations

Integrating the above results, the following qualitative model, illustrated in Fig. 5, emerges for the interactions of stress granules with the cytoskeleton. Upon stress, SG nucleation and growth occurs rapidly throughout the cytosol, and is complete within about 5 min from the appearance of the first SGs (Fig. 2). The dense f-actin network [37–39] of the lamellae confines stress granules, and its native retrograde flow expels them into the center of the cell. Subsequent contraction of actin network, now largely void of SGs, further guides SGs toward the cell center. Actin-driven transport thus concentrates SGs in the perinuclear region, increasing the rate of coalescence. Capillary interactions [24, 40–42] drive SGs toward regions of high microtubule density.

**FIG. 5.**
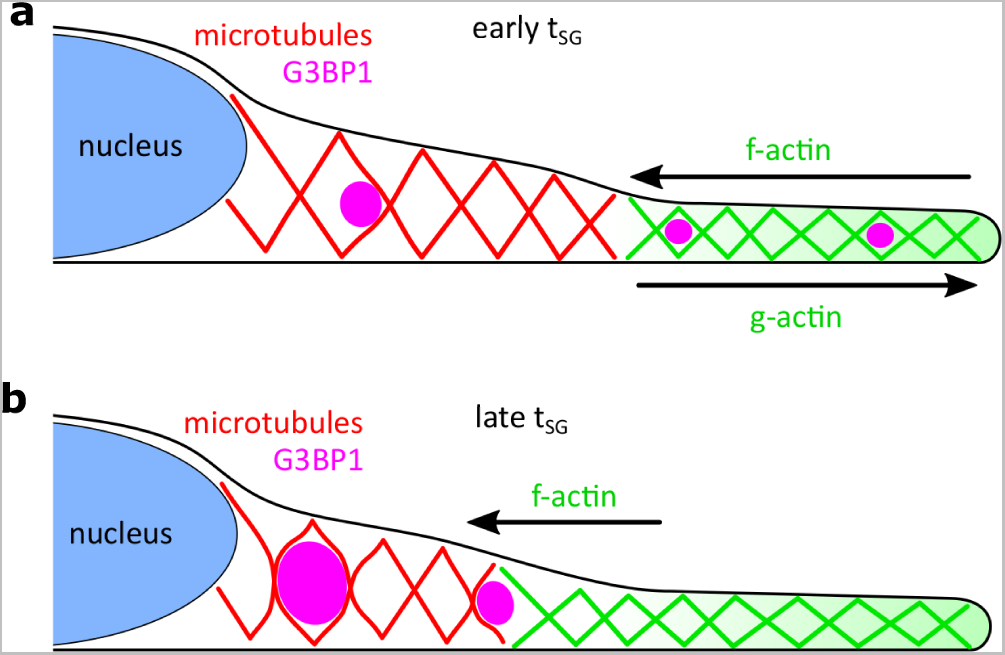
Model of the interactions between stress granules, actin and microtubules. **a**, right after nucleation and growth of SGs, SGs in the lamellar region are subject to retrograde flow and move towards the cell center, where they are handed over to the microtubule network. **b**, at longer times, actin contracts, leading to further inward flux of SGs. Favorable wetting interactions between microtubules and SGs lead to mutual deformations of SGs and the microtubule network, as well as stabile positioning of SGs in microtubule-rich perinuclear region.

To challenge this picture, we quantified the impact of various cytoskeletal perturbations on the spatial distributions of stress granules, and their structural correlations with actin and microtubules (see Materials and Methods V E).

When the microtubule network is de-polymerized with nocodozole [43], actin transport still drives SGs toward the center of the cell, Figure 6 **b**. However, in the absence of a microtuble-rich organizing center, SGs are more evenly distributed. Microtubule subunits, however, continue to be concentrated at the surface of stress granules, Figure 6 **f**. This is consistent with an adhesive capillary interaction between SGs and microtubule sub-units [24]. The full results for nocodazole-treated cells, including reference cells for all channels and correlation analyses, are shown in Supplemental Fig. 15.

**FIG. 6.**
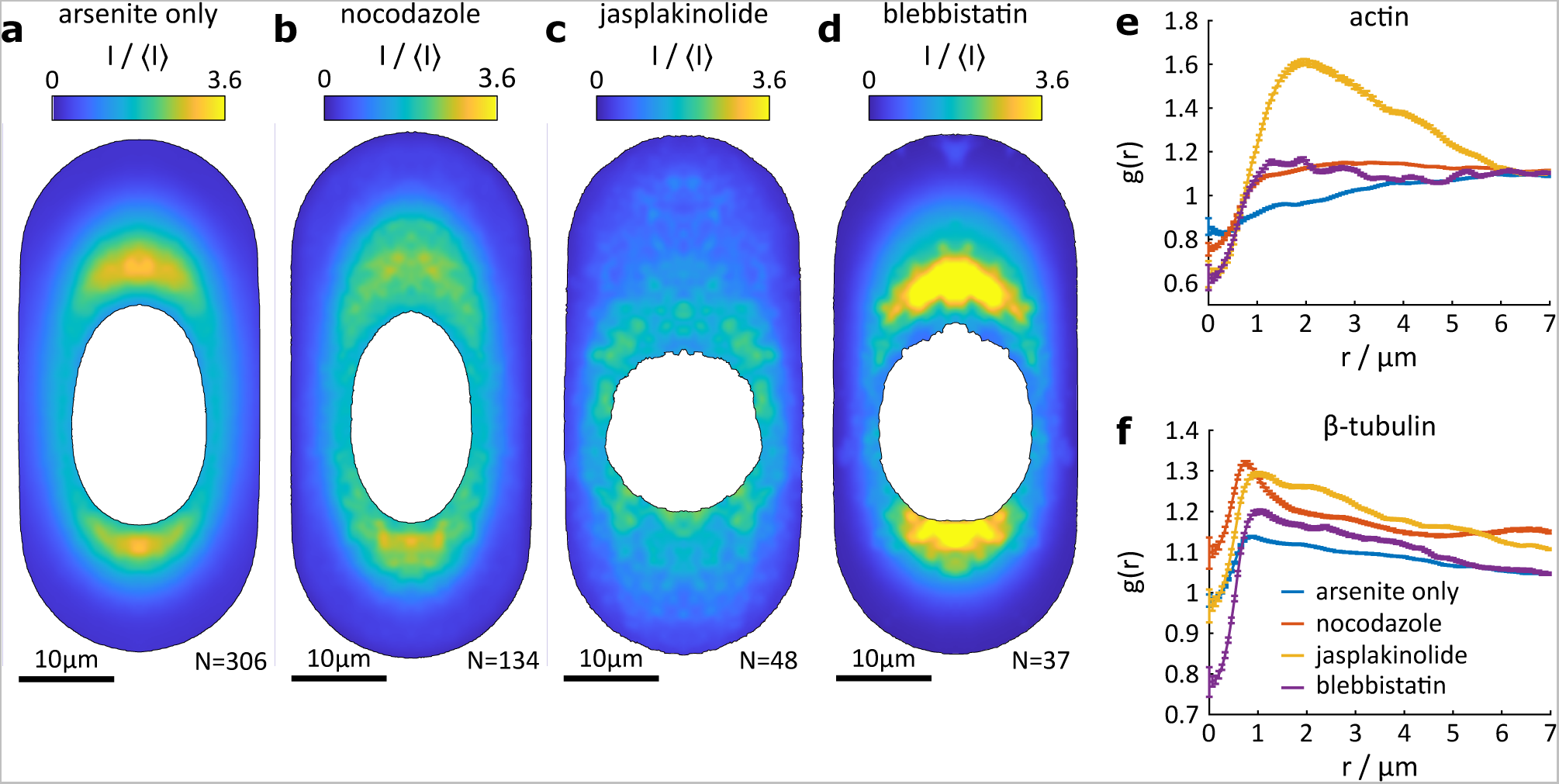
Perturbing the cytoskeleton affects the positioning of stress granules. Reference cells of G3BP1 for cells treated with arsenite only (**a**), treated with nodocazole before SG formation (**b**), treated with jasplakinolide before SG formation (**c**) and treated with blebbistatin before SG formation (**d**). Note that the colormap in panel **d** is saturated but kept identical for comparison purposes. An adjusted version can be found in Fig. 17 in the Supplement. *g*(*r*) of actin (**e**) and tubulin (**f**) for SGs with *R_SG_* = 0.65 µm for cells with different drug treatment.

Disruption of the actin network with jasplakinolide completely arrests actin dynamics and leads to aggregation of actin in amorphous masses around the cell nucleus [44, 45]. Live-cell experiments show that after nucleation and growth, SGs in jasplakinolide-treated cells do not migrate toward the center of the cell, and display reduced coalescence, see Supplementary Movie 4. As a result, SGs remain widely distributed throughout the cell, not just within the perinuclear region, see Fig. 6 **c**. Jasplakinolide seems to have a minimal effect on the interaction of SGs with microtubules (Fig. 6 **f**). On the other hand, SGs tend to be found near large clumps of actin (Fig. 6 **e**) The full results for jasplakinolide-treated cells are shown in Fig. 16 in the Supplement.

Disruption of myosin activity through treatment with blebbistatin has only a small effect on stress granules. As expected, blebbistatin treatment almost completely removes the stress-induced contraction of the actin network (see Fig. 17 **a** in the Supplement). Since retrograde flow is maintained at a reduced rate [34], SGs are still expelled from the lamellar region, and accumulate in micro-tubule rich regions of the cell center (Fig. 6 **d**). Blebbistatin does not have a major impact on the distribution of actin or microtubules around individual SGs (Fig. 6 **e, f**). The full results for blebbistatin-treated cells are shown in Fig. 17 in the Supplement.

Overall, these perturbations of the cytoskeleton are consistent with the emerging qualitative model proposed in Fig. 5.

## III. CONCLUSION

Stress granules nucleate, grow, coalescence, and coarsen in the presence of the active and heterogeneous networks of actin and microtubules. Overall, we find that the complex architecture and dynamic nature of the cytoskeleton gives rise to robust and reproducible positioning of stress granules through hydrodynamic coupling of granule motion to actin flow and adhesive capillary interactions with microtubules. Because of the generic nature of these physical interactions, we expect that they could also play a role in the dynamics of other membraneless organelles.

From a biological perspective, we expect that the final positioning of SGs in the perinuclear region may have functional importance. SGs are dynamic sites of mRNA regulation [9, 17, 46], suggesting a close interplay with the cell nucleus. Indeed, a number of regulatory RNA-binding proteins traffic between the nucleus and SGs [47–49]. Proximity of SGs to the nucleus might therefore ensure efficient signalling through short diffusion times of related proteins.

## IV. ACKNOWLEDGEMENTS

We thank Kathryn Rosowski for helpful discussions and assistance with cell culture.

## V. MATERIALS AND METHODS

For more detailed Materials and Methods, also refer to [24, 50].

### A. Cell culture

U2OS human osteosarcoma cells are grown in Dulbecco’s Modified Eagle Medium (DMEM, ThermoFischer, cat. num. 41966029) supplemented with 10 % fetal bovine serum (FBS, ThermoFischer, cat. num. 10270106), and 2 mM L-glutamine (ThermoFisher, cat. num. 25030024), at 37*^◦^*C in 5 % CO_2_. We use both wild-type cells, as well as U2OS RDG3. U2OS RDG3 have green fluorescent protein (GFP)-tagged G3BP1 to mark SGs as well as red flourescent protein (RFP)-tagged DCP1a, a processing body protein [27, 51, 52]. Note that G3BP1 does not co-precipitate with *β*−tubulin [19]. Both cell lines were kindly supplied by the Pelkmans Lab at the University of Zurich.

### B. Micropatterning

In order to ensure consistent cell shapes and cytoskeletal architecture between experiments, cells were plated on patterned coverslips. The optimal size of the pattern was determined during an initial trial where cells were placed on patterns of varying area. The final size of about 1240 µm^2^ was selected as the size where single cells adhered to one pattern and showed the best compliance to the pattern shape.

Patterned coverslips are fabricated as follows. 22 mm by 22 mm glass coverslips are first cleaned and rinsed with ethanol by hand and then further cleaned by exposure to deep UV in a UV/ozone cleaner (ProCleaner Plus BioForce Nanosciences) for 5 minutes. The clean coverslips are then incubated in 0.1 mg/ml poly-L-lysine-g-poly(ethyleneglycol) (PLL(20)- g[3.5]-PEG(2), SuSoS AG) for one hour. A quartz photomask (printed by Deltamasks) containing transparent features in an otherwise opaque chrome coating is cleaned in the same fashion as above with ethanol and 5 min in the UV/ozone cleaner. The transparent features are 25 µm-by-30 µm rectangles with two semi-circular caps of radius 12.5 µm at the ends, similar to the patterns used by Oakes et al. [28]. One coverslip contains several hundred features. Features are spaced at least 100 µm apart from edge to edge in all directions. The coated coverslips are mounted onto the photomask with the coated side in contact with the mask, using the capillary forces of a thin water film between glass and mask (about 8 µl water per coverslip). The mounted coverslips are then exposed to deep UV light coated-side up in the UV/Ozone cleaner through a chrome on quartz photomask (printed by Deltamasks) for ten minutes. After UV treatment, coverslips are detached by floating the coverslips with water and stored in distilled water for up to three weeks.

During experiment preparation, the patterned coverslips are first submerged in 70% ethanol for at least 5 minutes and subsequently rinsed three times with phosphate-buffered saline (PBS). Next, the coverslips are coated with 20 µg/ml of fibronectin (Sigma-Aldrich) for 12 minutes. For fixed cell exerpiments, the coated coverslips are rinsed with PBS three times and transferred to a 6-well plate with one coverslip per well. Cells are plated at a concentration of about 5 · 10^4^ cells per well 4–6 hours prior to experiments to ensure sufficient spreading. Longer wait times typically lead to an increasing sub-population of cells that divide on the pattern.

### c. Fixed cell experiments

#### 1. Immunofluorescence

Cells are plated on patterned coverslips as outlined above. After specified durations of treatment with 0.5 mM sodium arsenite (Sigma-Aldrich) to induce stress granule formation [25] or additional drugs treatment depending on the experiment, U2OS cells are fixed with 4 % formaldehyde for 15 min, permeabilized with 0.1 % Triton X-100 in PBS for 20 min, and blocked with 5 mg/ml bovine serum albumin (Sigma-Aldrich) for one hour. Fixed cells are incubated with primary antibodies overnight at 4*^◦^*C. Primary antibodies were mouse anti-G3BP (1:500 in blocking solution, abcam ab56574) and rabbit anti-β-tubulin (1:200 in blocking solution, abcam ab6046). Note that G3BP does not co-precipitate in β-tubulin immunoprecipitation and is commonly used as a stress granule marker [19, 52, 53]. On the next day, cells are stained in solution containing secondary antibodies for 1 hour at room temperature. The secondary antibodies are Rhodamin Red-X anti-mouse IgG (1:500 in blocking solution, Jackson ImmunoResearch, cat. num. 115-295-003) and Alexa 647 anti-rabbit IgG (1:500 in blocking solution, Jackson ImmunoResearch), cat. num. 111-605-144). Actin is stained using 488 nm-phalloidin (ThermoFischer, cat. num. A12379). DNA is stained by incubation in DAPI solution (1:1000 in PBS, SigmaAldrich, cat. num. D9542-10MG) for 15 min at room temperature. Coverslips are finally mounted in ProLong Gold or Dimond (both Thermo Fisher). Cells are washed three times with PBS in between different steps.

#### 2. Fixed cell imaging

Confocal stacks of fixed cells are imaged on a Nikon Ti2 Eclipse with Yokogawa CSU-W1 spinning disk and 3i 3iL35 Laser Stack using a 100x oil objective with a numerical aperture of 1.45 and a Hamamatsu Orca-flash 4.0 camera. The spatial resolution in the focal (xy-)plane is 0.065 µm/pixel and the step height (z-direction) is 0.2 µm.

The theoretical diffraction limit for the longest used wavelength (640 nm), here defined as the full width half maximum of the theoretical point spread function of a confocal microscope, is 160 nm, calculated as with numerical aperture NA [54]. For the actual optical setup, we assume a diffraction limit of three pixels (corresponding to 195 nm).

#### 3. Cell Detection and Sorting

Image analysis is carried out using matlab and is largely automated to allow for efficient and reproducible processing of a large number of cells. The protocol outlined below is for fixed cells but the data processing of live-cell experiments is largely analogous. Each set of confocal data acquired on the microscope contains one cell. The cell is identified in the xy-plane using the *regionprops* function on an overlay of the maximum projections along z of all channels as well as a wide-field image of the nucleus through the DAPI stain. For the accurate detection of the cell shape in the focal plane, the image is sharpen before thresholding. All images are cropped around the detected cell in the xy-plane. To determine the z-coordinate of the base of each cell (cell base height) we calculate the sum of the median and 80th-percentile for each xy-plane of the cytoskeleton channel. With only a small rim of background around the cropped cells, the median of the image is slightly below the median intensity of the cell and serves as a proxy for the overall structure of the cell. The 80th-percentile captures more pronounced features, such as individual filaments, while being robust against outliers. The cell base height is then determined as the z-coordinate corresponding to the maximal gradient of this intensity measure, which reliably detects the side of the cell attached to the coverslip. We assume an error of this measurement of ±1 z-step. The maximum intensity z-slice is then typically 2 to 4 slices above the cell base height. Cells in which this distance falls out of this interval are discarded. Moreover, cells with an area outside 85 to 110 % of the area of the prescribed pattern (1257 µm^2^) are discarded. Cells with a ratio of the short principal axis to the long principal axis outside the interval [0.43 0.55] are also discarded. The corresponding ratio of the pattern itself is 0.45. These criteria have been chosen based on the corresponding histograms of all cells to discard outliers.

The background intensity of each channel is approximated from the intensity in the corners of the cropped image outside the detected cell. Each channel is corrected for this background intensity individually.

Multiple cells are recorded on each coverslip in one acquisition session. Within one such a batch, all patterns have the same orientation. The orientation of the patterns is determined as the mean orientation of cells from one batch. Batches with less than 10 cells remaining after filtering the cell area and shape are discarded. Individual cells with an orientation that deviates by more than 3*^◦^* relative to the pattern orientation are also discarded. All cells are then rotated by the mean orientation of the pattern to ensure consistent cell orientation across samples. The remaining cells are aligned such that the centroid of each cell is in the center of the xy-plane. Because the nucleus is not always exactly in the center of the cell, cells are arranged such that the centroid of the nucleus always falls in the same hemisphere of the image, i.e. some cells are rotated by 180*^◦^*.

#### 4. Intensity Normalization

The intensities of the G3BP1, actin and β-tubulin channels of each cell are individually normalized by their respective mean intensity ⟨I⟩ to account for differences in protein expression or staining. In order to define this mean intensity, we select a set of representative pixels. The x- and y-coordinates of representative pixels are those that fall inside the cell shape but outside the cell nucleus, based on the maximum projection along the z-coordinate of all channels. Pixels outside the cell or inside the nucleus were set to *NaN* (not a number) throughout all following analyses. The z-position of the representative pixels are chosen to be the second to fourth z-coordinate above the detected cell base height. Typically, the third z-coordinate above the cell base is the maximum intensity z-plane. The mean intensity for a given channel is then calculated as the mean across all representative pixels. Each channel is then normalized by the respective mean intensity in that cell.

#### 5. Stress granule detection

SGs are detected in confocal stacks of the G3BP1 channel. We determined the background by blurring the image stack with a box-kernel of a size larger than typical granules. This background is subtracted from the confocal stack. The background-adjusted image stack is then blurred with a Gaussian kernel with σ = 1 pixel in the xy−plane to decrease shot noise. The resulting filtered stack is then re-scaled such that intensity values are positive by adding the minimum intensity value in a stack to each voxel.

Next, 101 threshold values evenly spaced between the maximal mean intensity value within any z-slice of the image stack and the global maximum intensity are probed. For each threshold, entities above threshold are detected using the matlab function *regionprops3*, recording the position, volume, voxel list, mean and median intensity for each connected group of voxels. With these information, we assign a quality factor to each threshold. The quality factor is defined as the sum of the median intensity and the mean intensity across all detected particles for a given threshold divided by the threshold intensity. The median intensity as a function of threshold level is smoothed using a running average to decrease noise. By definition, this factor is larger or equal to two. This quality factor will be large if a blob is brighter than background. Too low thresholds are penalized as they pick up the surrounding of a bright region of pixels yielding low mean and median intensity, too high thresholds are penalized by dividing by the threshold. Consequently, this measure approaches a value of two for too low as well as too high thresholds. The threshold maximizing this quality factor is taken. Cells where the global maximum is not well defined against other local maxima, i.e. here at least 7.5 % higher, are discarded. All detected granules are subject to further tests. In fixed cell data, all detected entities are inflated by dilation with a sphere of three pixels radius. Particles that fuse upon dilation are discarded to ensure a minimal distance between granules. Further, SGs where the voxels, by which the granule grew upon dilation, are brighter than 55 % of the intensity of the detected granule are also discarded to ensure sufficient contrast against the local environment. In the fixed cell data, granules, typically very small, that reside below or above the cell nucleus were discarded as they fall outside the cell mask.

We define the granule radius based on the maximum gradient of the intensity profile across the SG interface, see [24, 50]. SGs where the maximal gradient across the interface is further away than 2 pixels from edge of the SG as defined by the threshold are discarded. Granules with a radius below the optical resolution limit are discarded, see Methods section V C 2.

Aside from the characteristics (position, orientation, volume, etc.) a number of images are saved for each granule. All images are centered on the centroid of a given granule and show the xy-plane closest to the granule centroid. Note that pixels within these images that fall outside the cell mask are set to *NaN*. Note that the mask for fixed cells also excludes the nucleus, while the mask in live cell data only defines the cell edge. The size of extracted images is about 14x14 µm, sufficient to capture also the long range deformations of the cytoskeleton around granules.

#### 6. Construction of the reference cell

The alignment of all cells in three dimensions allows to construct cell stacks by overlaying all cells that belong to the same experimental conditions, e.g. duration of arsenite treatment. Each cell stack is blurred in the xy-plane using a Gaussian kernel with a variance of 4 pixels. While this method retains the overall intensity, noise as well as single filaments are blurred to suppress short range fluctuations. To capture the finite size of the cytosol as well as of the nucleus, each cell is masked by the cell outline and the nucleus shape. Based on these stacks, the reference intensity for any spatial coordinate is calculated as the mean intensity across all cells at this position, omitting the data from cells where the given pixel is masked. If more than half of the intensity values for a given coordinate fall within these masks of the nucleus or cell outline, the pixel is considered outside the cytoplasm of the reference cell. This way, the reference cell also serves as a mask for the expected cell shape.

The symmetry along the short axis has deliberately been broken by rotating all cells such that the cell nucleus falls into the same half of the cell. The remaining symmetry of the pattern allows to fold the cell along the center line of the long axis to enhance the statistics for the calculation of the reference cell. Using N cells, this process yields 2N intensity values for all spatial coordinates in one half of the cell. The full reference cell is then recovered by mirroring the result along the center line.

Note that, due to fluctuations in the height of the cells, we only considered the intensity values between the cell base height up to 1.2 µm (6 slices) into the cell as reliable. Stress granules outside this range are not considered in the following analysis.

#### 7. Correlation analysis

Correlation analysis has been used in the cell, e.g. to study protein distributions in the lipid membrane [55, 56]. The main difference here is, that we account for the finite cell shape and heterogeneous cell architecture, see also [24, 50].

We calculate distribution maps (e.g. Fig.3) in the same way as averaged images only that each individual image is now normalized by the corresponding image of the reference cell. Normalization is done by point-wise division with the respective image originating from the reference cell of that channel at the same location. Note that, if a pixel is masked in either reference or G3BP1 and β-tubulin image, it is considered masked and set to NaN. Consequently, different pixels of a distribution map may not have the same number of pixels that contributed to the calculation of the corresponding intensity value.

Fluctuations in the number of contributing data sets for different pixel locations introduces a non-linearity when further analyzing distribution maps, which has to be taken into account when calculating the radial distribution g(r) of a distribution map. To capture the varying statistical weight, we do not take the radial average of the distribution map in question. Rather we take the average over all intensity values from the individually reference-cell-normalized images that the distribution map is calculated from at distance r from the center, again omitting masked entries. This way we know how many data points contribute to each entry in g(r). The error of g(r) at a given distance r is then calculated as the standard error, i.e. the standard deviation of the contributing values divided by the square root of the number of contributing values.

The surface-relative distribution function g*_s_*(d) is calculated from the reference-cell-normalized images for each granule individually, as it is the basis of the calculation of the partitioning coefficients for each granule. Here, the distance d for a given pixel location is calculated as the minimal distance to the outline of the stress granule. The outline itself has a value of d = 0 and pixels inside the granule have negative entries. The outline is defined by the stress granule detection routine. g*_s_*(d) is then calculated by averaging the intensities of pixels with the same d, omitting masked entries. The error is again calculated as the standard error.

To validate the distribution maps, we perform a negative control where we extract the images corresponding to a detected SG not from the cell in which the granule was detected, but from a different cell that has been exposed to the same conditions. If all cells are listed in a table, we essentially shift the entries in this cell listing by one position, i.e. matching SGs with a different cell. This method yields random, yet biologically plausible, input that is normalized by the exact same segment of the reference cell. These negative control distribution maps, or the corresponding g(r) should ideally assume a uniform value of one.

#### 8. Calculation of stress granule volume fraction

To determine the probability of finding SGs in regions of a certain actin or *β*−tubulin intensity in the cell, we first assign an intensity value I/⟨I⟩ to each SG that is determined by the intensity of the voxel in the three-dimensional reference cell of either actin or tubulin that corresponds to the centroid of each granule. The volume fraction of SGs within regions of a reference cell *ϕ_SG_* is then calculated as the sum over the number of voxels in i SGs with a given I/⟨I⟩ (N*_SG,i_*) normalized by the respective number of voxels of the reference cell with this value of I/⟨I⟩ (N*_V_*) and normalized by the number of SGs N

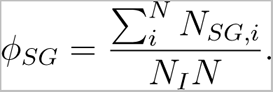

Note that also N*_i_* is calculated based on the three-dimensional information of the reference cells. This calculation of *ϕ_SG_* is based on the volumetric extend of each SG as determined by the SG detection routine. These values, however, typically overestimates granule volume because the initially detected SG radius R*_SG_* does not align with the maximum gradient in G3BP1 across the interface. In g*_s_*(d), the maximum gradient is typically at a distance of minus one to two pixel into the granule, see e.g. Fig. **?? c**. While R*_SG_* can readily be corrected to align with the position of the maximum gradient in intensity of G3BP1 across the granule interface, correcting N*_SG_* accordingly is not so easy due to several caveats, such as a non-cubic size of voxels due to differences in resolution in xy and z. To estimate the error of the volume fraction, we employ a second method to calculate *ϕ_SG_*. Here, we consider the corrected R*_SG_* in the xy-plane and assume a spherical granule shape:

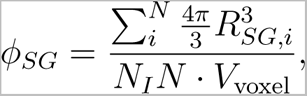

with volume per voxel V_voxel_. This method assumes a spherical shape and thus likely underestimates the granule volume in many instances. Together, we have two calculation modes that should over- and underestimate *ϕ_SG_* for SGs with regular shapes. Indeed, *ϕ_SG_* calculated based on the corrected R*_SG_* typically yields values below *ϕ_SG_* based on the SG detection. For irregular shapes, i.e. SGs that break symmetry in one plane, e.g. SGs with an oblate shape mainly extending in the xy-plane, *ϕ_SG_* assuming a spherical shape may overestimate the actual volume. Such irregular shapes are more pronounced for larger SGs, typically found in regions of high tubulin intensity. Taken together, we expect that *ϕ_SG_* based on the initial SG detection overestimates granule volume but is fairly robust against shape changes as the actual volumetric extent is considered. *ϕ_SG_* calculated using the corrected R*_SG_* and assuming a spherical shape is expected to under-estimate *ϕ_SG_* as long as SG shapes do not break symmetry in three dimensions. A comparison of the values of *ϕ_SG_* yielded by these methods for a given I/⟨I⟩ thus provides an estimate of the actual volume fraction.

### D. Live cell experiments

Live cell experiments are done using U2OS RDG3 cells, while fixed cell experiments are done with U2OS wild-type cells. The protocols, however, are identical for either cell type.

Cells are plated on patterned coverslips as outlined above. Instead of a 6-well plate, however, coverslips are mounted in a magnetic chamber that can be placed onto the microscope (Live Cell Instruments, Chamlide chambers CM-S22-4).

In case live stains are used to stain the cytoskeleton, the cell media is switched to media containing the appropriate concentration of the stains at least 2 hours before the start of experiments and about 1–2 hours after plating the cells on the patterns. All live-cell experiments presented here were done using the recommended concentration for SPY-tubulin and SPY-FastAct cytoskeletal tags (Spirochrome). Note that SPY actin stains only bind f-actin. Further, SPY actin is conjugated to a ligand from a jasplakinolide derivative, a drug that stabilized f-actin and inhibits network turnover [39]. Similarly, SPY tubulin is conjugated to the ligand docetaxel, which can also cause adverse effects in the microtubule network. Both ligands are modulated to minimize cell toxicity of the probes [57]. At the recommended concentration, Spirochrome claims no adverse effects in the cell.

#### 1. Live cell imaging

Cells are imaged at 37*^◦^*C and 5 % CO_2_ in normal cell media. Imaging is done on a Nikon Ti2 Eclipse microscope with cage incubator and environmental chamber (both okolabs), using a 100x oil objective with numerical aperture of 1.49. Data are recorded using a Yokogawa CSU W1 spinning disc with 50 µm pinhole-size, together with a laser bench (Oxxius) including a 405 nm line at 180 mW, 488 nm and 561 nm at 200 mW and a 640 nm line at 300 mW and a Photomerics Prime 95B camera. Together with the 100x objective this gives a spatial resolution in the xy-plane of 0.11 µm per pixel. Note that the resolution limit is the practically the same as on the setup used to image fixed cells. Due to the larger pixel size, however, we consider features with a size of 2 pixels (220 nm) optically resolved. Cells are imaged using Nikon’s perfect focus system (PFS) to keep the cell in the focal plane.

The laser intensity was adjusted during trials to find a balance between induced phototoxicity and signal strength for each channel. Typically, 6 confocal stacks are recorded per cell at a spacing of 1 µm at 30 sec intervals. Cells are treated with 150 µM arsenite to induce SGs. This is done by adding 3 µl of 50 mM arsenite to 1 ml of media in the chamlide chamber and subsequent careful stirring by pipetting several hundred µl of media up and down. A given experiment is started as soon as arsenite is added to the chamber. Note that this arsenite concentration is lower than the concentration used in fixed cell experiments (500 µM). This concentration has been determined throughout various trials to induce SGs in most cells while ensuring that cells withstand the stress for more than one hour. While the typical arsenite concentration in literature is also 500 µM, it is not uncommon to use lower concentration, e.g. [58]. The lower concentration suggests, however, that live-cell imaging, especially with a confocal microscope, is not free of artifacts and may induce further cell stress upon imaging.

Data from cells that exhibit odd behavior, including presence of SGs in the absence of arsenite, abnormal changes in morphology or detachment from the substrate, are discarded. Especially temperatures in the incubation chamber on the microscope above 37*^◦^*C are observed to induce SGs. Such data is also discarded. In case SGs exhibit unusual behavior, such as granulation or apparent hardening, at later times of arsenite treatment, the experimental data is cut multiple minutes before the onset of such changes in SG behavior to ensure consistency between experiments.

#### 2. Live cell data analysis

Cells are analyzed using matlab code derived from the code used for fixed cells. The cell detection routine is modified such that the cell is tracked over time to account for drift in the xy-plane. We approximate the error of the drift correction over time to be ± one pixel. Cells are rotated such that the nucleus, or rather where we expect the nucleus to be based on the images of the actin and G3BP1 channel, falls onto the same side of the cell.

#### 3. Stress granule detection

SGs are detected using the same routine as outlined above (Methods and Materials V C 5). We do, however, not perform the filtering step that discards SGs that are very close to each other in order to register SG coalescence and fusion. Also, the local contrast requirement is relaxed, considering granules where the surrounding is up to 70% as bright as the SG itself, to enhance detection of small and dim SGs.

The calculation of SG volume is based on the SG detection by thresholding. The detected SG outline may deviate from the position of the maximum gradient in fluorescence intensity across the granule interface that is otherwise used to define the granule radius. The difference between threshold-detected granule outline and maximal gradient position is, however, below one pixel. For the exemplary cell (Fig. 1 and Fig. 2 **a** - **c**), we find a mean difference between the maximum of the intensity gradient and the threshold-detected interfaces of -0.27 pixel (0.018 µm), i.e. we may slightly overestimate the SG volume.

#### 4. Reference construction

Reference cells are constructed by local averaging. For live-cell data, we do not have as rich information in the z−direction as in fixed cells as we only record data within the cell to minimize phototoxicity. Consequently, it is difficult to ensure good alignment in z and to accurately define the bottom of the cell. Reference cells for the live-cell data are therefore not resolved in z but calculated by averaging of the maximum projections along z of the confocal stacks of a given cell at a given time. The time axis is aligned between cells such that t*_SG_* = 0 min corresponds to the point in time when the first SGs were detected in a given cell. Pixels where more than half of the data from contributing cells have non-*NaN* entries are considered part of the reference cell, analogous to the construction of the reference cells for fixed cells. Note that the intensity normalization is not as precise as in fixed cells because the extent of the cell nucleus is not known, i.e. we cannot accurately define the cytosol, just the inside of the cell. The intensity of a given maximum projection of a given channel of a given cell at time t*_SG_* is instead normalized by the average intensity across the maximum projection considering only the inside of the cell.

#### 5. Particle image velocimetry

We performed particle image velocimetry (PIV) on the maximum projections of the actin and G3BP1 channel for each cell to extract the flow fields of actin and G3BP1 over time. PIV is done in matlab using code calling on functions from the matlab plug-in *PIVlab* [59]. The smallest interrogation window has a size of 8 by 8 pixel, i.e. 0.9 by 0.9 µm. To make use of the different cells we recorded, we average PIV results from individual cells, inspired by previous works [60]. For the averaging of the PIV results across different cells, each cell is masked by the respective cell outline, where pixels outside the cell are set to *NaN*. Further, we introduce a minimal contrast criterion to ensure reliability of the correlation analysis. For each interrogation window, we calculate the standard deviation of the local intensity. Only those windows are accepted that have a standard deviation 1.75 (4) times larger than the standard deviation in the background for the actin (G3BP1) channel. Windows that do not exceed this contrast measure are set to NaN. PIV results are further averaged by collecting data over time, binned by steps of 2.5 min. This method yields stacks of PIV results of up to 175 individual measurements. Finally, the PIV results are averaged across all measurements in a given window across cells and time to arrive at *PIV ensembles*. Windows in the PIV ensembles where less than 25% of the measurements contribute a non-*NaN* value were discarded. Note, that the error in cell detection to correct for drift may cause errors in the PIV results. We estimate errors from the cell detection to be ±1 pixel. Assuming that the error is symmetric, however, we expect that the performed averaging over many cells and time steps minimizes this error. We perform a similar analysis based on tracking of SGs, yielding consistent flow speeds, see Supplementary Fig. 13.

#### 6. Tracking of stress granules

Tracking of SGs is done using an established tracking function in the lab. The code considers the x−, y− and z−coordinates, as well as R*_SG_*, to identify the same SGs across frames. This code is based on the work of John C. Crocker from the University of Chicago and has been updated over time, by Eric R. Dufresne and others (see online repositories https://site.physics.georgetown.edu/matlab/).

### E. Drug treatments

Cells are treated with nocodazole (Sigma-Aldrich, CAS 31430-18-9) at a concentration of 1.67 µM, jasplakinolide (Sigma-Aldrich, CAS 102396-24-7) at 50 nM and blebbistatin (Sigma-Aldrich, CAS 856925-71-8) at 20 µM for 30 min before adding arsenite to the media in order to ensure that the desired effect on the cytoskeleton is present in the cells at the onset of the stress response. The timing of 30 min was determined through live-cell trials using SPY-FastAct and SPY-tubulin.

## Supplementary Material

### VI. SUPPLEMENTAL FIGURES

**FIG. 7.**
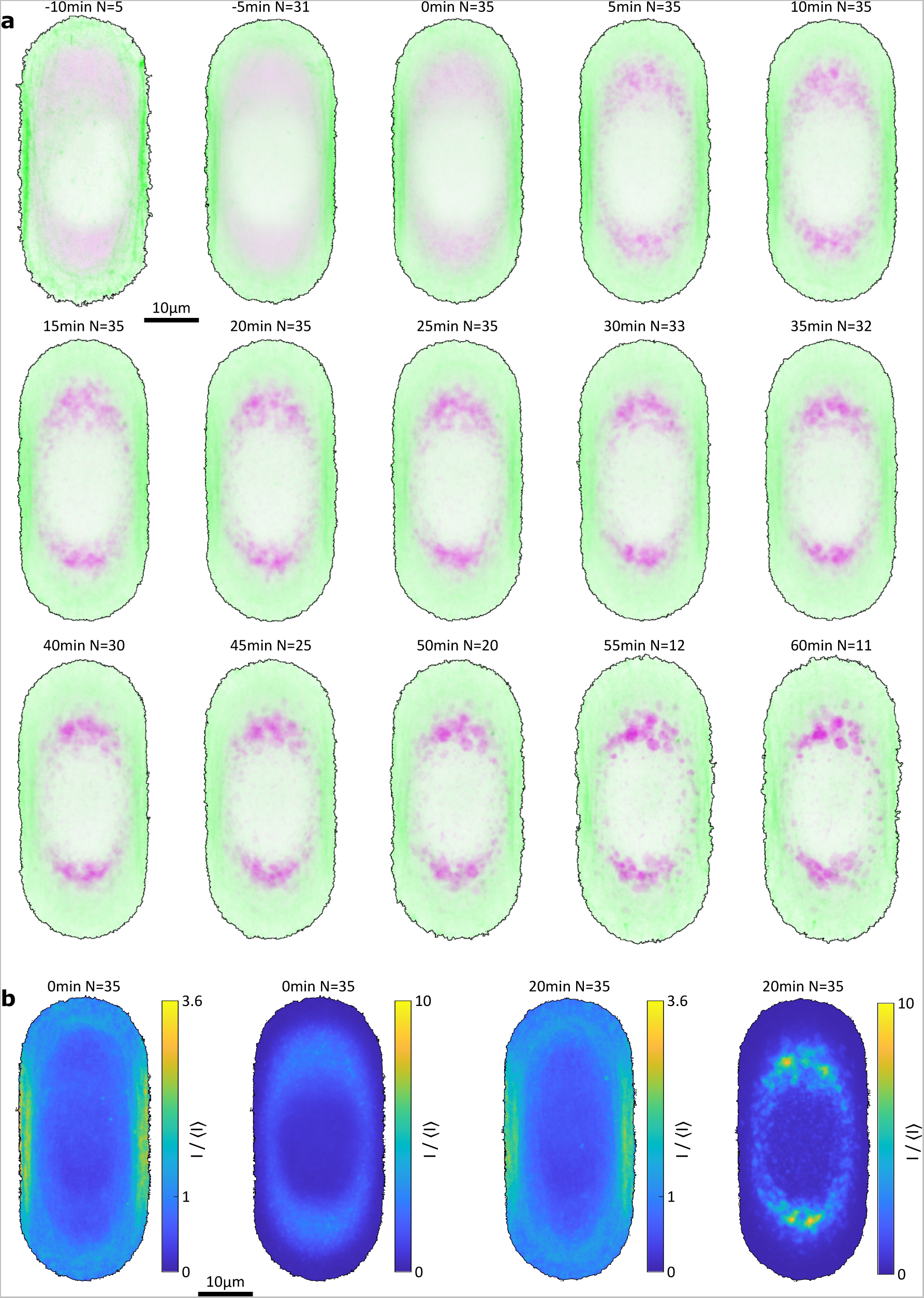
Reference cells for actin and G3BP1 over time extracted from live-cell data. **a**, time-resolved reference cell averaged over maximum projections from *N* cells for actin (green) and G3BP1 (magenta). *t* = 0 min corresponds to the point in time when the first SG has been detected in cells. **b**, reference cells for 0 min and 20 min with the full scale bar.

**FIG. 8.**
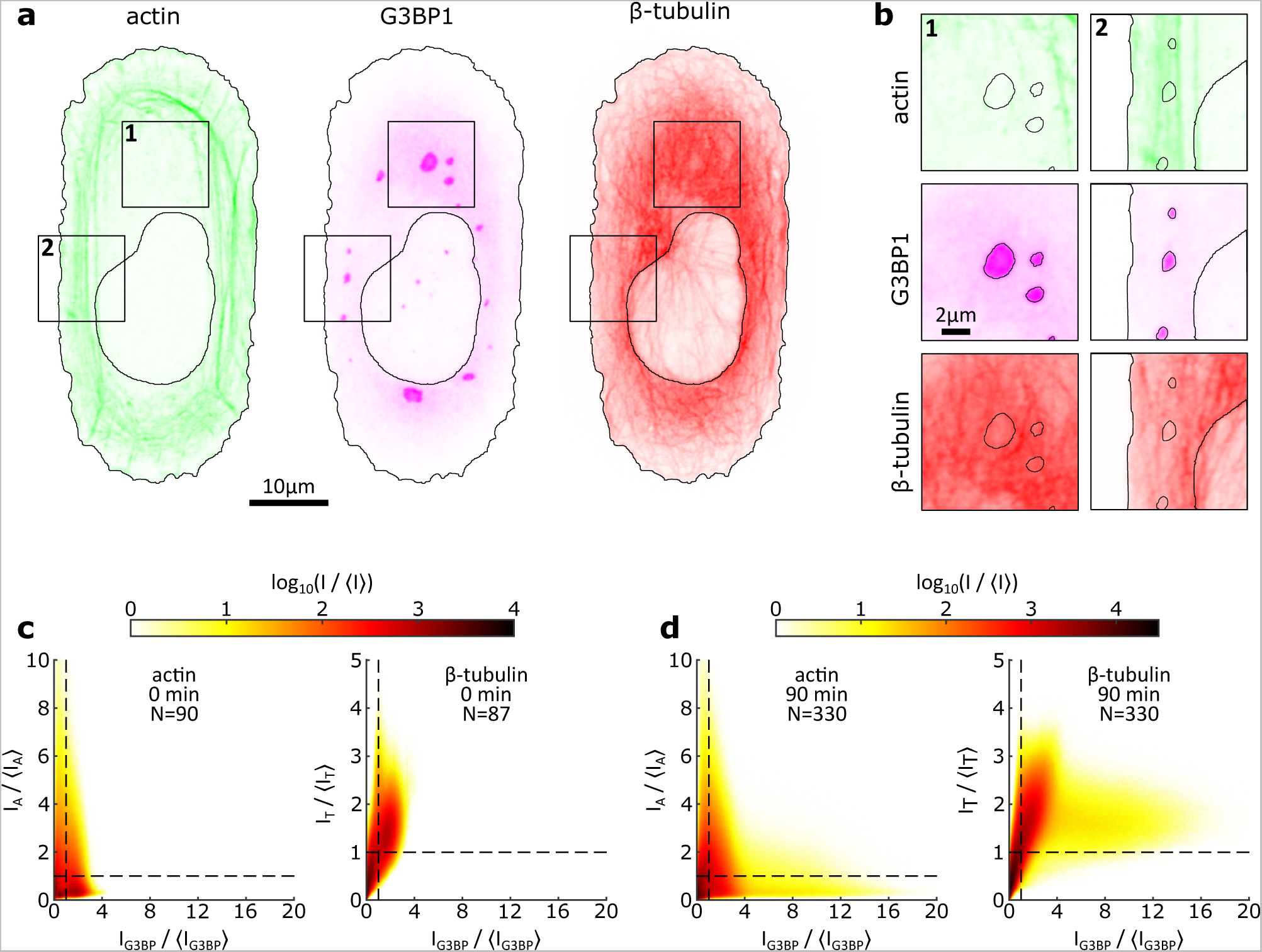
Interactions of stress granules with the cytoskeleton. **a**, exemplary xy-slices of the actin, G3BP1 and *β*-tubulin channels of the same cell after 90 minutes of arsenite treatment. The black line shows the outline of the detected cell and nucleus. **b**, zoomed images centered on individual granules as indicated by the squares in **a**. The images are overlaid with the outline of detected granules. **c** and **d**, spatial correlation of G3BP1 with actin and tubulin before (**c**) and after (**d**) arsenite treatment, i.e. at 0 min and 90 min treatment respectively. The correlation is calculated across one *xy*-slice taken from each cell with a total of *N* contributing to the analysis. All data are normalized by the mean intensity of the given channel in the respective cell ⟨*I*⟩.

**FIG. 9.**
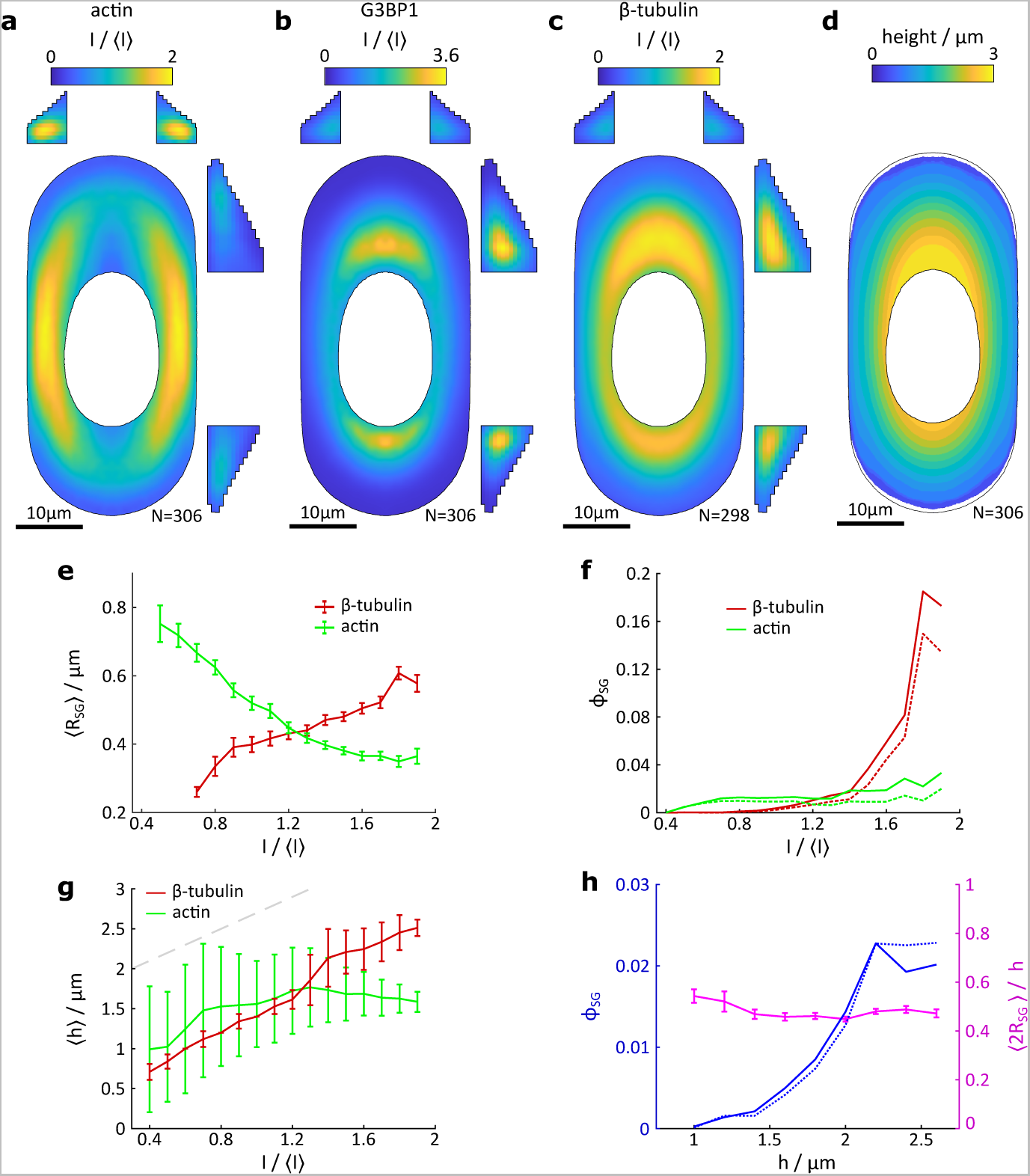
Stress granules are correlated with low actin intensity and high tubulin intensity but also with cell height. **a**, reference cell for actin after 90 min of arsenite treatment. *N* gives the number of contributing cells. The side views shown are taken along the respective center line across the cell. The side views are stretched three fold in the *z*−direction compared to the scale bar in the *xy*−plane and are overlain with the approximated cell height. **b** and **c**, the reference cells for G3BP1 and *β*−tubulin. **d**, height map of the cell. Voxels are considered inside the cell if the respective value of the tubulin reference cell is greater than 0.4. The black outline in the *xy*-plane shows the extent of the reference cells and almost aligns with the extent of the cell height map using this measure. **e**, the average SG radius ⟨*R_SG_*⟩ as a function of the value of the respective reference cell at the centroid of each granule. Error bars are given as standard error. **f**, volume fraction *ϕ_SG_* given as the probability that a voxel with a given intensity in the actin or *β*−tubulin reference cell is part of the volume of a SG. Solid lines indicate *ϕ_SG_* calculated using the number of registered voxels of SGs, dashed lines show *ϕ_SG_* calculated based on the radius of each granule given by the location of the maximum gradient in the G3BP1 channel. **g**, average cell height ⟨*h*⟩ as a function of *I/*⟨*I*⟩ for actin and *β*−tubulin. Errors bars show standard error. The grey dashed line shows a slope of one. **h**, *ϕ_SG_* as a function of cell height. Solid and dashed lines indicate the different calculation method as before. The rig_2_h_9_t *y*-axis shows the average SG diameter, ⟨2*R_SG_*⟩, compared to the local cell height.

**FIG. 10.**
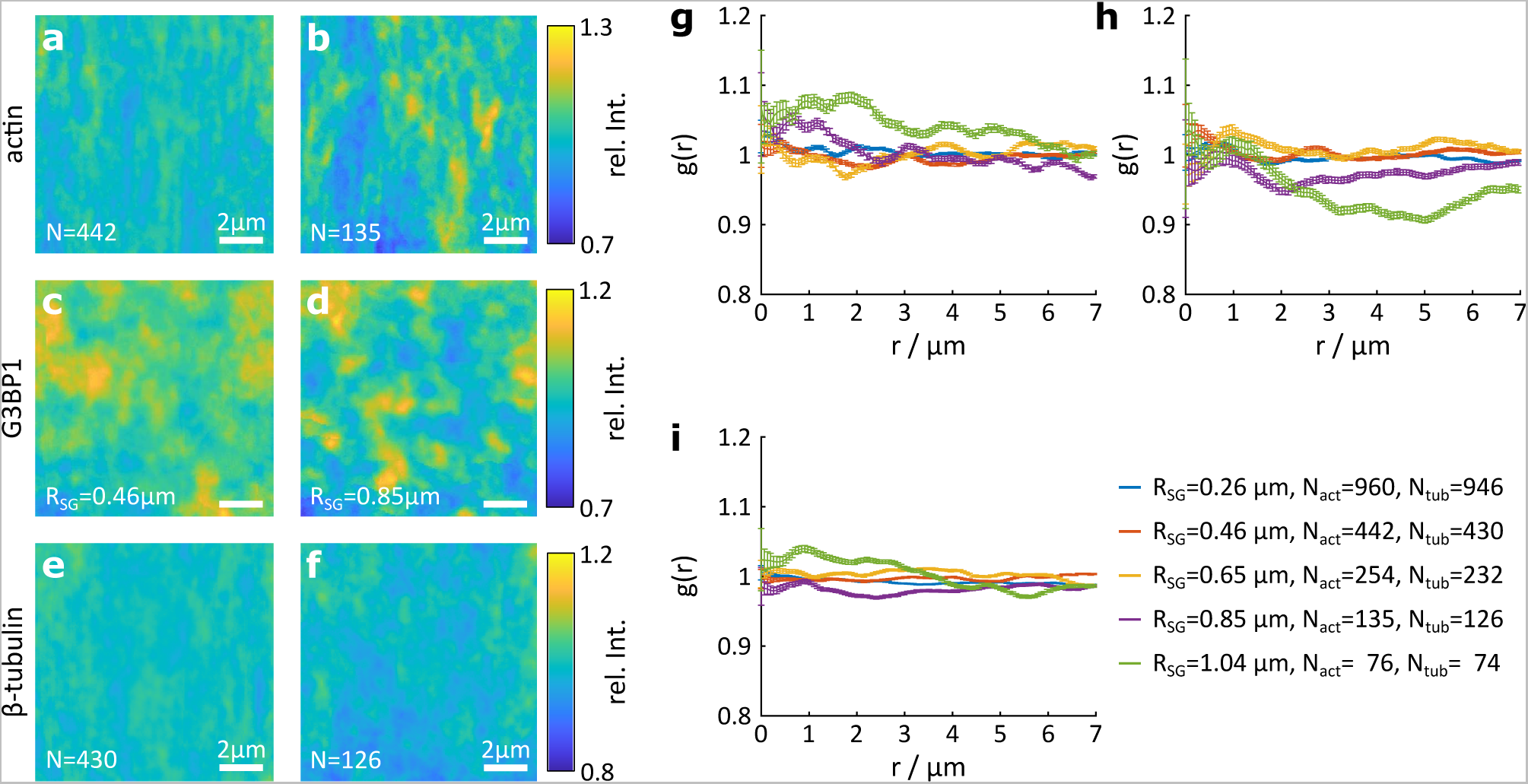
Negative control for distribution maps and *g*(*r*) at 90 min of arsenite treatment, see Fig. 3 **e** - **m**. **a**, **b**, negative control for the distribution maps for actin for granules with *R_SG_* in the interval of [0.39 µm 0.52 µm] and [0.78 µm 0.91 µm] respectively. *N* gives the number of contributing SGs. **c**, **d**, the corresponding controls for G3BP1. The number of contributing SG is identical to those for actin. **e**, **f**, control for *β*−tubulin around SGs of the same size. Note that the number of contributing SGs is slightly different. **g**, *g*(*r*) of the control for actin around SGs of varying *R_SG_*. **h**, *g*(*r*) of the control for G3BP1 around SGs of varying *R_SG_*. **i**, *g*(*r*) of the control for *β*−tubulin around SGs of varying *R_SG_*.

**FIG. 11.**
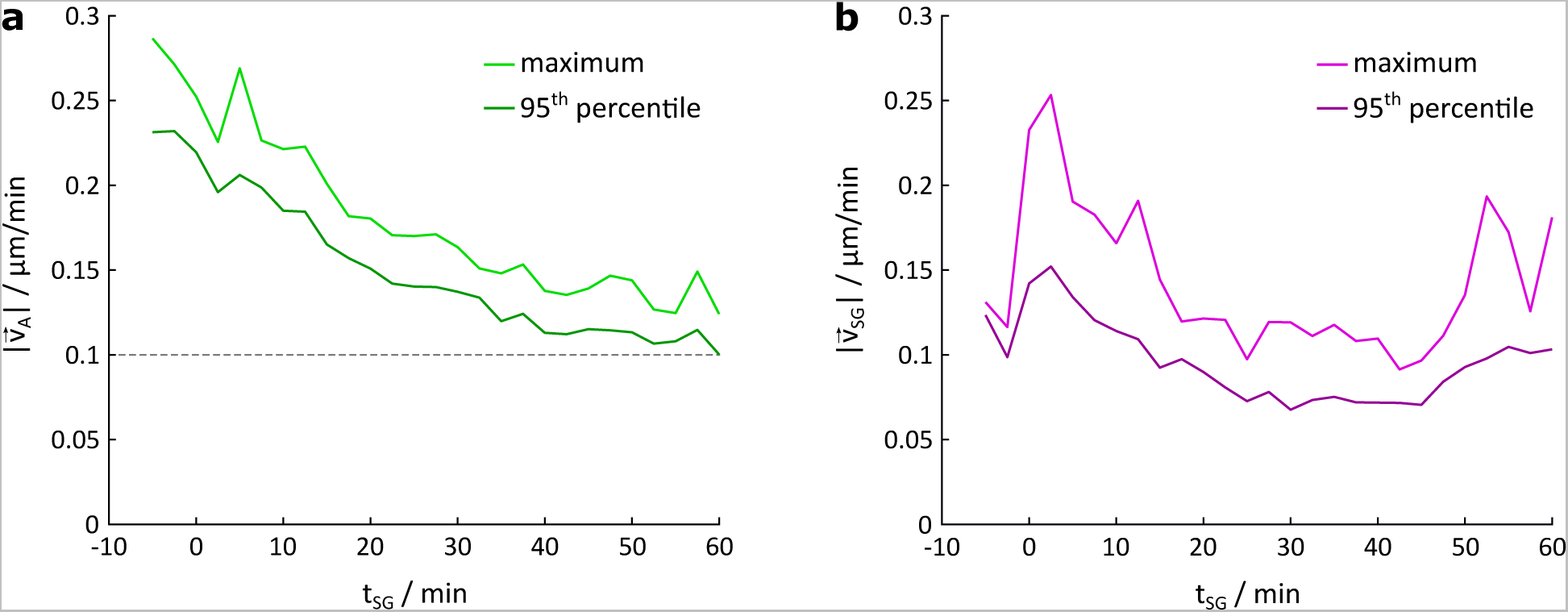
Maximum and 95^th^ percentile of the flow speeds found throughout the PIV ensemble at a given *t_SG_* for actin (**a**) and G3BP1 (**b**). The value of 0.1 µm/min is a guide to the eye to compare to the flow speed of actin contraction (see Fig. 12).

**FIG. 12.**
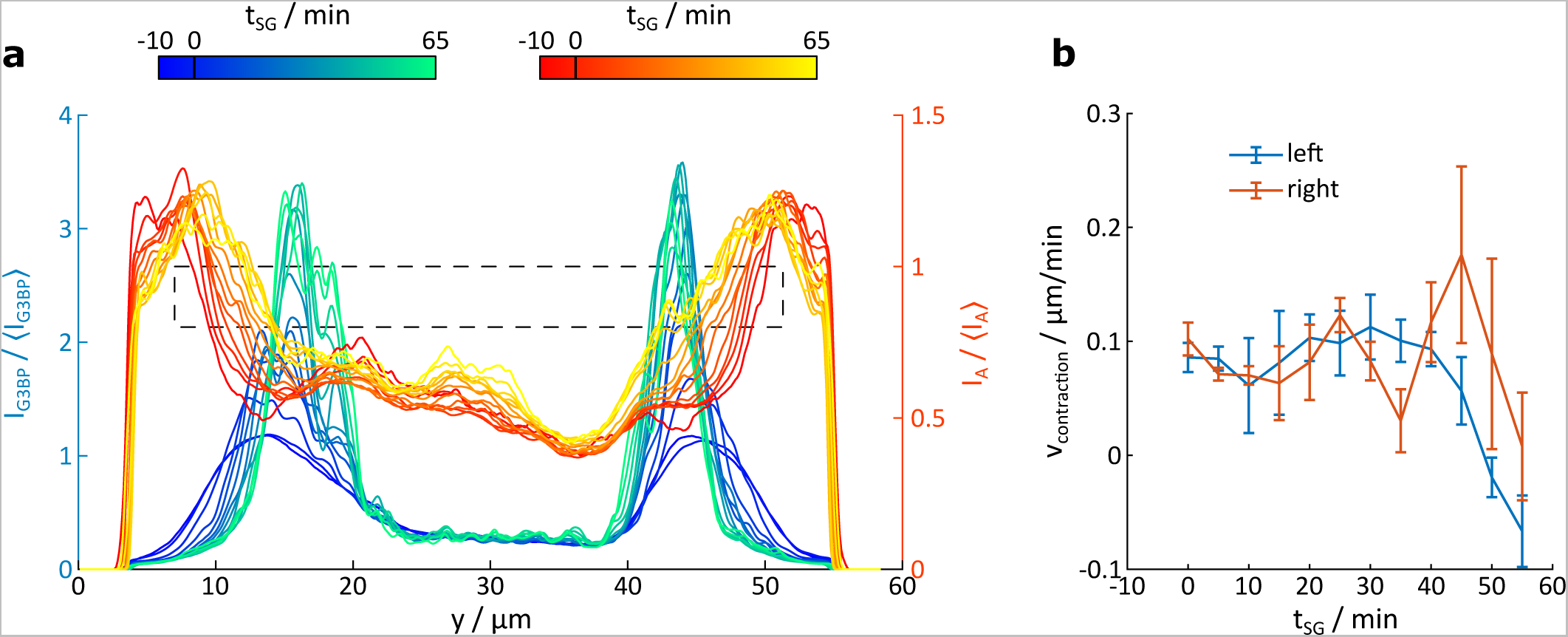
The actin network and SGs move towards the cell center over time. **a**, intensity across the long axis of symmetry of the time-resolved reference cells for the G3BP1 (blue-green) and actin channel (red-yellow). Each line shows the average over 5 min. *t_SG_* = 0 min corresponds to the time when first SGs form. **b**, average contraction speed of actin at the left and right side of the cell calculated by averaging the actin positions over time within the dashed box indicated in panel **a**.

**FIG. 13.**
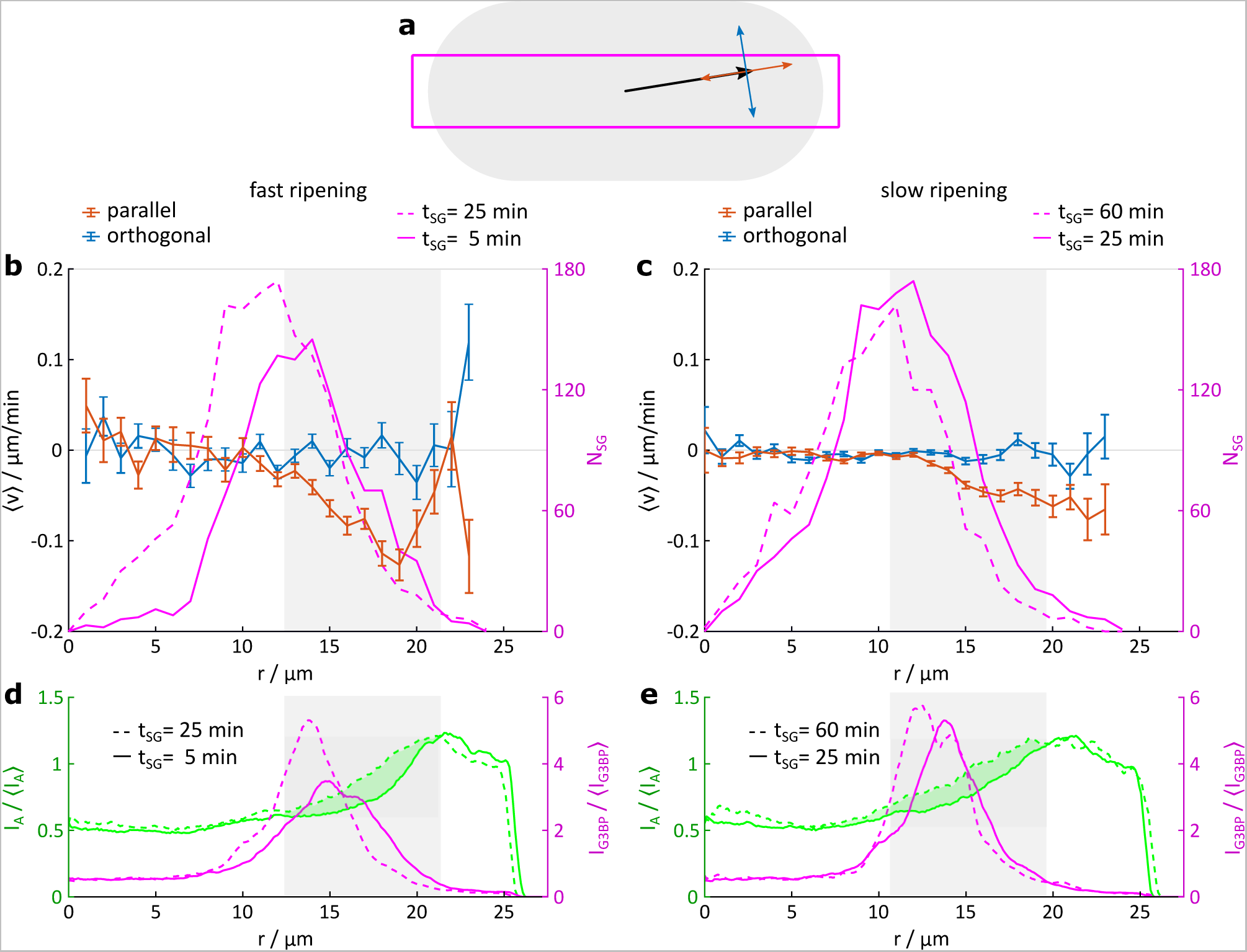
Granule tracking reveals that directional SG flow occurs in regions that are subject to contraction of the actin network. **a**, sketch of the construction. Only SGs within a 14 µm wide band along the long axis of the cell (outlined in grey) are considered, i.e. within the magenta box. The speed vector of each SG within the box is decomposed into a component orthogonal and parallel to the local position vector originating from the cell center (black arrow). **b**, average speed ⟨*v*⟩ orthogonal and parallel to a radial vector from the cell center as a function of distance from the center *r* within the first 25 min after first SGs have been detected. *N_SG_* gives the sum of considered SGs in two time bins at *t_SG_* = 5 and 25 min. **c**, analogous ⟨*v*⟩ collected over data with *t_SG_ >* 25 min. *N_SG_* shown for *t_SG_* = 25 and 60 min. Note that the center of the cell is largely filled with the cell nucleus that extends roughly 10 µm from the center. **d**, intensity profiles of the time resolved reference cells calculated over the same interrogation area sketched in **a** for *t_SG_* = 5 and 25 min for the actin and G3BP1 channels. The area shaded in grey highlights the region over which the actin network moves inward between 5 to 25 min. **e**, corresponding intensity profiles of the time resolved reference cells for *t_SG_* = 25 and 60 min for both channels. The area shaded in grey highlights the region over which the actin network moves inward between 25 to 60 min.

**FIG. 14.**
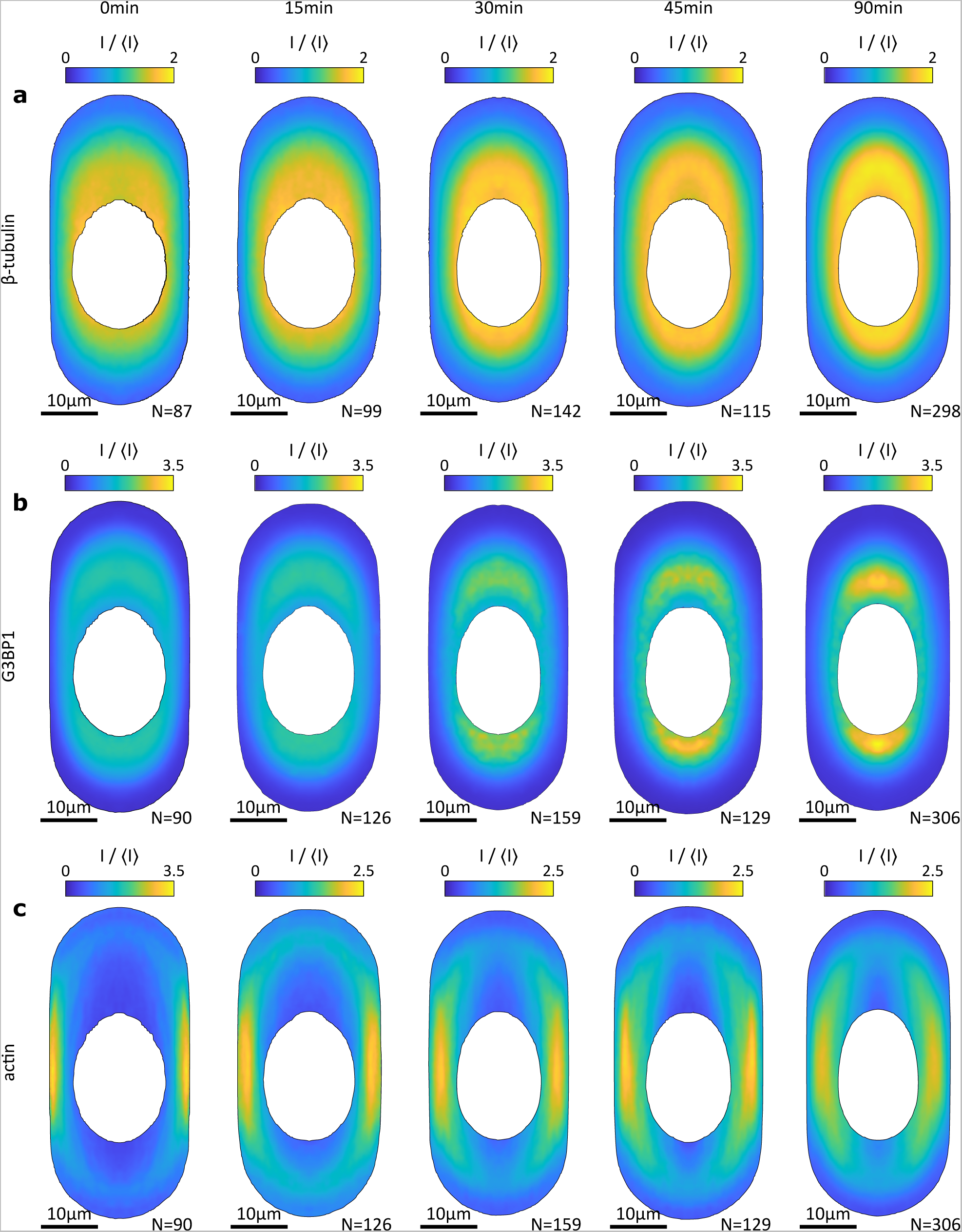
Reference cells for, **a**, *β*−tubulin, **b** G3BP1 and, **c**, actin without arsenite treatment (0 min), and after 15, 30 45 and 90 min of arsenite treatment. *N* gives the number of cells contributing to each reference cell. Note that the color map for actin without arsenite treatment is not the same as for other cells.

**FIG. 15.**
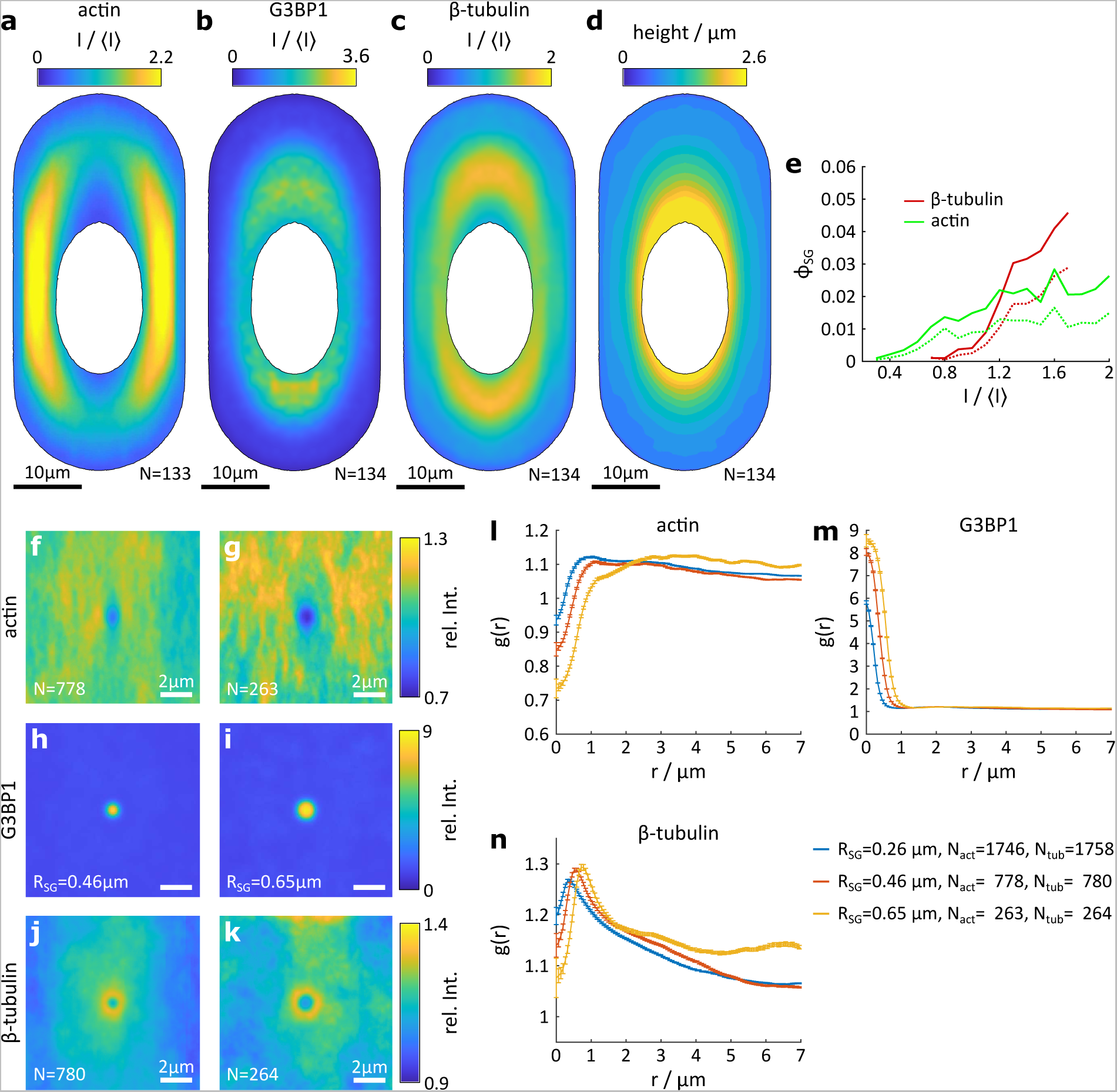
Interactions of SGs with the cytoskeleton in nocodazole-treated cells. **a**, **b**, **c**, reference cell of the actin, G3BP1 and microtubule channels averaged over *N* cells. The intensity is normalized by the average intensity in each individual cell and channel ⟨*I*⟩. The black line indicates the extent of the cytosol. **d**, height profile of the reference cell. **e**, volume fraction *ϕ_SG_* of stress granules in regions of the cell with a given intensity of the actin and *β*−tubulin channel. Solid lines indicate *ϕ_SG_* calculated using the number of registered voxels of SGs, dashed lines show *ϕ_SG_* calculated based on the radius of each granule given by the location of the maximum gradient in the G3BP1 channel. **f**, **g**, distribution map of actin around stress granules of roughly spherical shape (principal axis ratio below 1.5) and with a radius in the interval from 0.39 to 0.52 µm (indicated as *R_SG_* = 0.46 µm) and from 0.78 0.91 µm (indicated as *R_SG_* = 0.85 µm). *N* gives the number of contributing SGs. **h**, **i**, distribution map of G3BP1 for small and larger SGs. **j**, **k**, distribution map of *β*−tubulin for small and larger SGs. **l**, radial distribution function *g*(*r*) of actin for granules of varying size. **m**, **n**, *g*(*r*) for G3BP1 and *β*−tubulin respectively. The control for the data in panel **f** - **n** is shown in

**FIG. 16.**
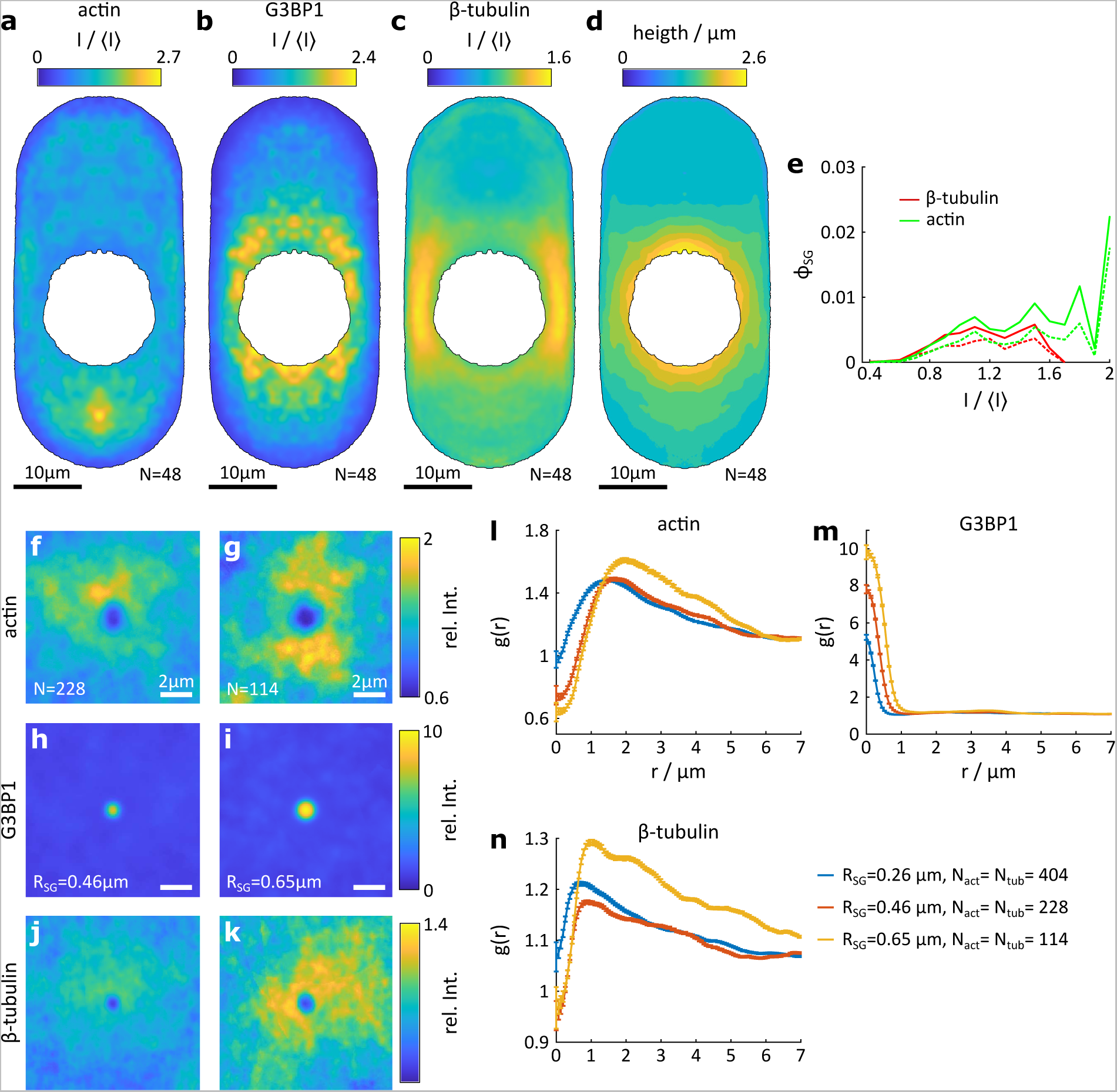
Interactions of SGs with the cytoskeleton in jasplakinolide-treated cells. **a**, **b**, **c**, reference cell of the actin, G3BP1 and microtubule channels averaged over *N* cells. The intensity is normalized by the average intensity in each individual cell and channel ⟨*I*⟩. The black line indicates the extent of the cytosol. **d**, height profile of the reference cell. **e**, volume fraction *ϕ_SG_* of stress granules in regions of the cell with a given intensity of the actin and *β*−tubulin channel. Solid lines indicate *ϕ_SG_* calculated using the number of registered voxels of SGs, dashed lines show *ϕ_SG_* calculated based on the radius of each granule given by the location of the maximum gradient in the G3BP1 channel. **f**, **g**, distribution map of actin around stress granules of roughly spherical shape (principal axis ratio below 1.5) and with a radius in the interval from 0.39 to 0.52 µm (indicated as *R_SG_* = 0.46 µm) and from 0.78 0.91 µm (indicated as *R_SG_* = 0.85 µm). *N* gives the number of contributing SGs. **h**, **i**, distribution map of G3BP1 for small and larger SGs. **j**, **k**, distribution map of *β*−tubulin for small and larger SGs. **l**, radial distribution function *g*(*r*) of actin for granules of varying size. **m**, **n**, *g*(*r*) for G3BP1 and *β*−tubulin respectively. The control for the data in panel

**FIG. 17.**
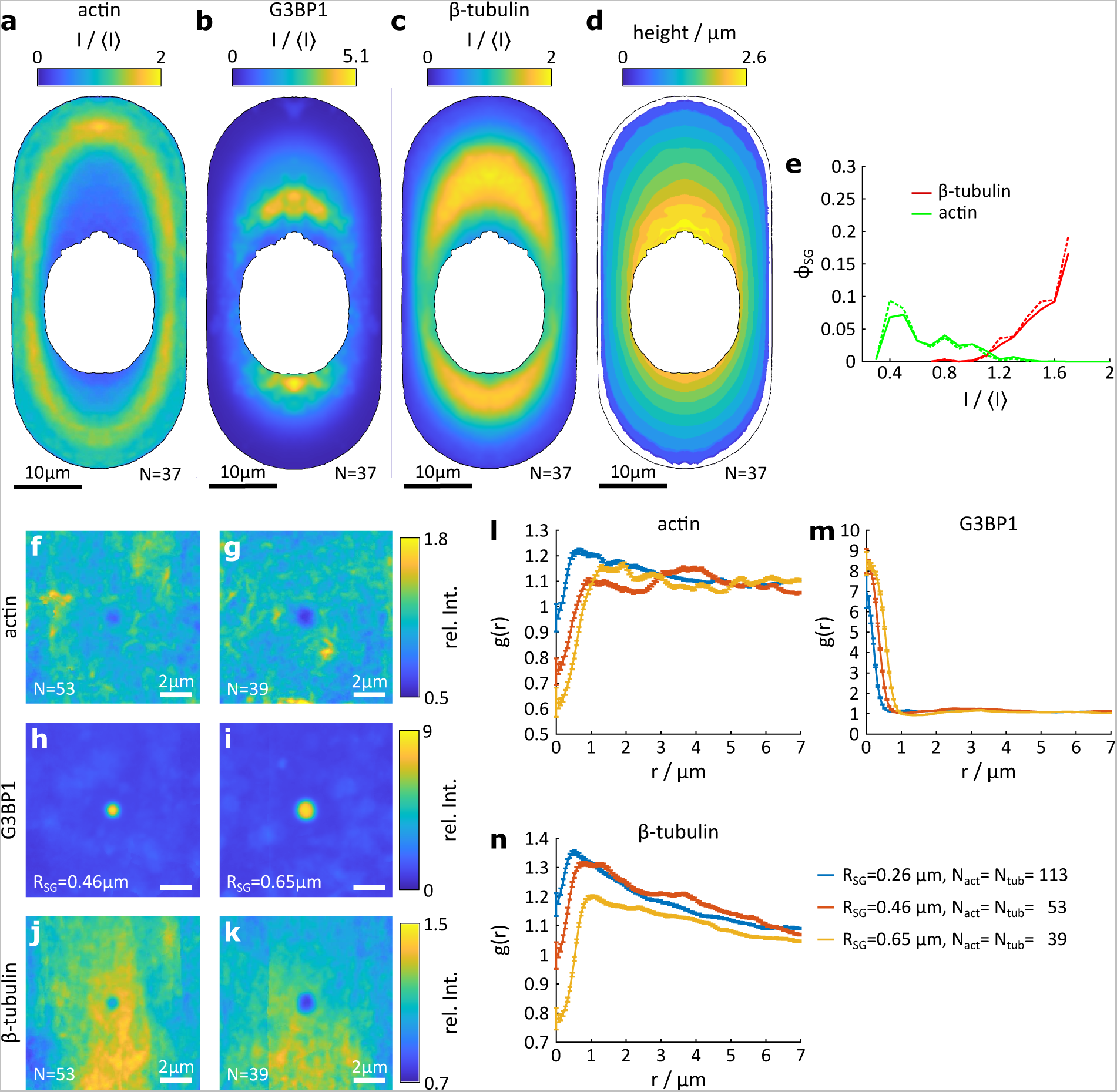
Interactions of SGs with the cytoskeleton in blebbistatin-treated cells. **a**, **b**, **c**, reference cell of the actin, G3BP1 and microtubule channels averaged over *N* cells. The intensity is normalized by the average intensity in each individual cell and channel ⟨*I*⟩. The black line indicates the extent of the cytosol. **d**, height profile of the reference cell. **e**, volume fraction *ϕ_SG_* of stress granules in regions of the cell with a given intensity of the actin and *β*−tubulin channel. Solid lines indicate *ϕ_SG_* calculated using the number of registered voxels of SGs, dashed lines show *ϕ_SG_* calculated based on the radius of each granule given by the location of the maximum gradient in the G3BP1 channel. **f**, **g**, distribution map of actin around stress granules of roughly spherical shape (principal axis ratio below 1.5) and with a radius in the interval from 0.39 to 0.52 µm (indicated as *R_SG_* = 0.46 µm) and from 0.78 0.91 µm (indicated as *R_SG_* = 0.85 µm). *N* gives the number of contributing SGs. **h**, **i**, distribution map of G3BP1 for small and larger SGs. **j**, **k**, distribution map of *β*−tubulin for small and larger SGs. **l**, radial distribution function *g*(*r*) of actin for granules of varying size. **m**, **n**, *g*(*r*) for G3BP1 and *β*−tubulin respectively. The control for the data in panel **f** - **n** is shown in Fig. 19 in the Supplement. ^36^

**FIG. 18.**
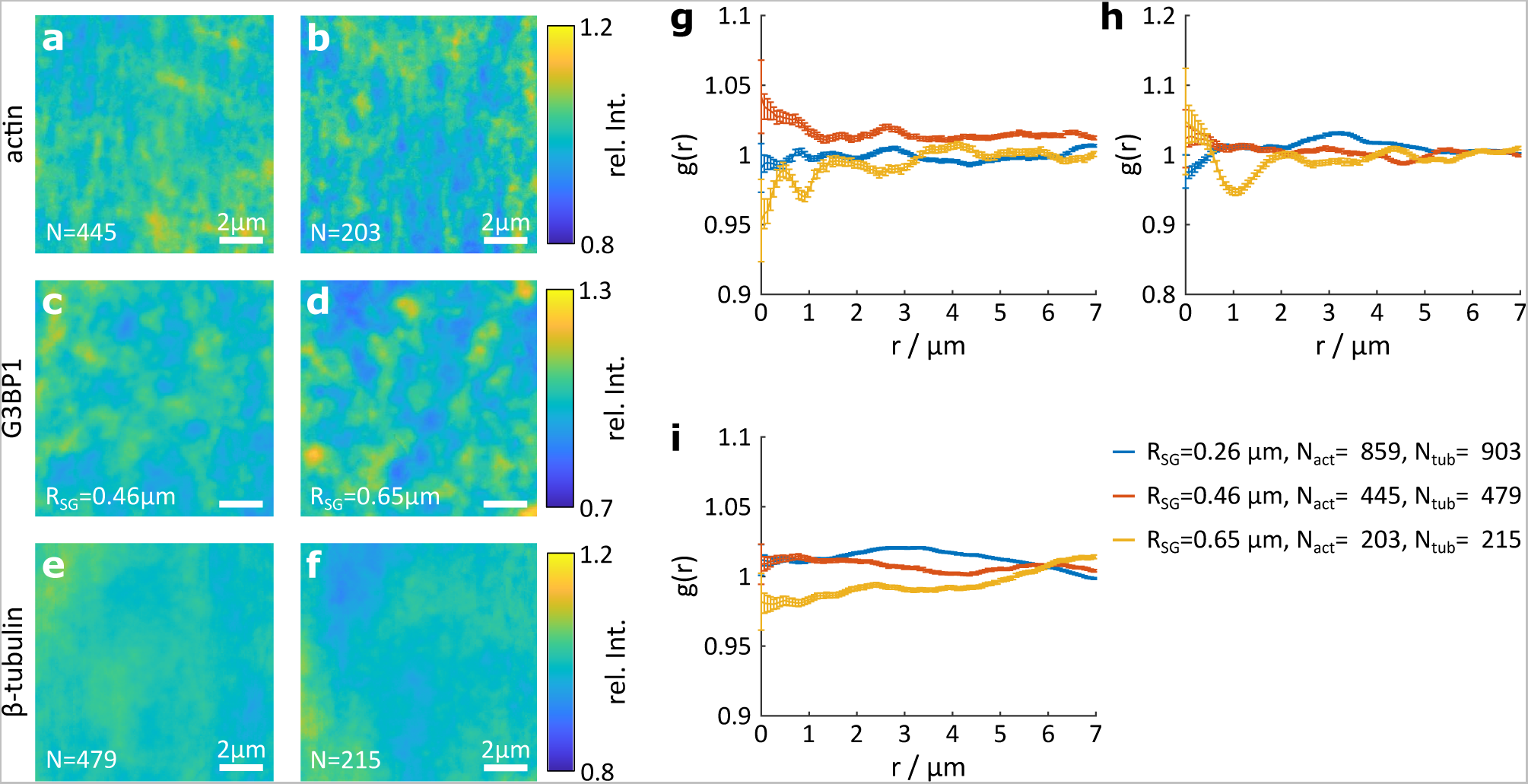
Negative control for distribution maps and *g*(*r*) after 30 min of nocodazole treatment before 90 min of arsenite treatment, see Fig. 15. **a**, **b**, negative control for the distribution maps for actin for granules with *R_SG_* in the interval of [0.39 µm 0.52 µm] and [0.59 µm 0.72 µm] respectively. *N* gives the number of contributing SGs. **c**, **d**, the corresponding controls for G3BP1. The number of contributing SG is identical to those for actin. **e**, **f**, control for *β*−tubulin around SGs of the same size. Note that the number of contributing SGs is slightly different. **g**, *g*(*r*) of the control for actin around SGs of varying *R_SG_*. **h**, *g*(*r*) of the control for G3BP1 around SGs of varying *R_SG_*. **i**, *g*(*r*) of the control for *β*−tubulin around SGs of varying *R_SG_*.

**FIG. 19.**
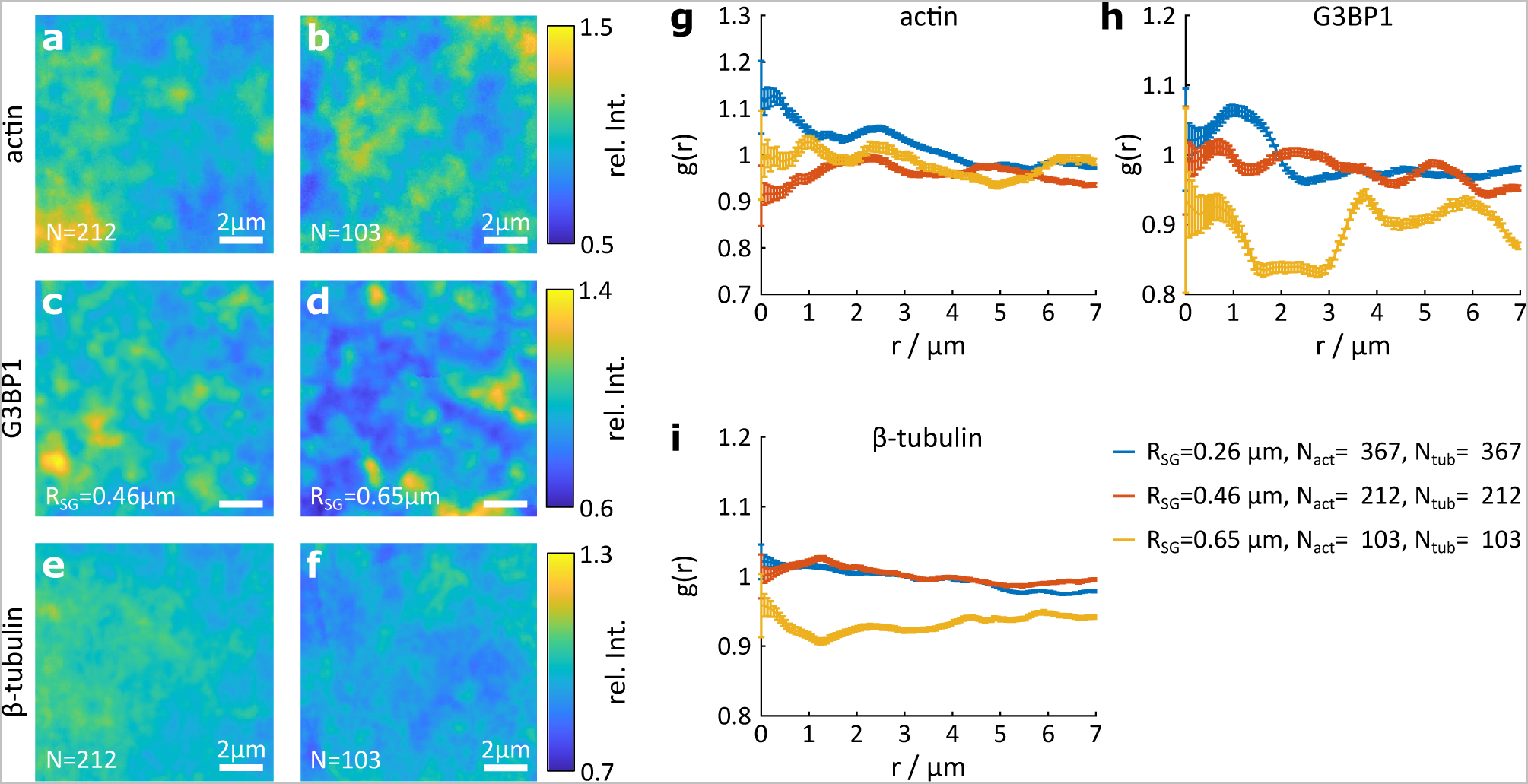
Negative control for distribution maps and *g*(*r*) after 30 min of jasplakinolide treatment before 90 min of arsenite treatment, see Fig. 16. **a**, **b**, negative control for the distribution maps for actin for granules with *R_SG_* in the interval of [0.39 µm 0.52 µm] and [0.59 µm 0.72 µm] respectively. *N* gives the number of contributing SGs. **c**, **d**, the corresponding controls for G3BP1. The number of contributing SG is identical to those for actin. **e**, **f**, control for *β*−tubulin around SGs of the same size. Note that the number of contributing SGs is slightly different. **g**, *g*(*r*) of the control for actin around SGs of varying *R_SG_*. **h**, *g*(*r*) of the control for G3BP1 around SGs of varying *R_SG_*. **i**, *g*(*r*) of the control for *β*−tubulin around SGs of varying *R_SG_*.

**FIG. 20.**
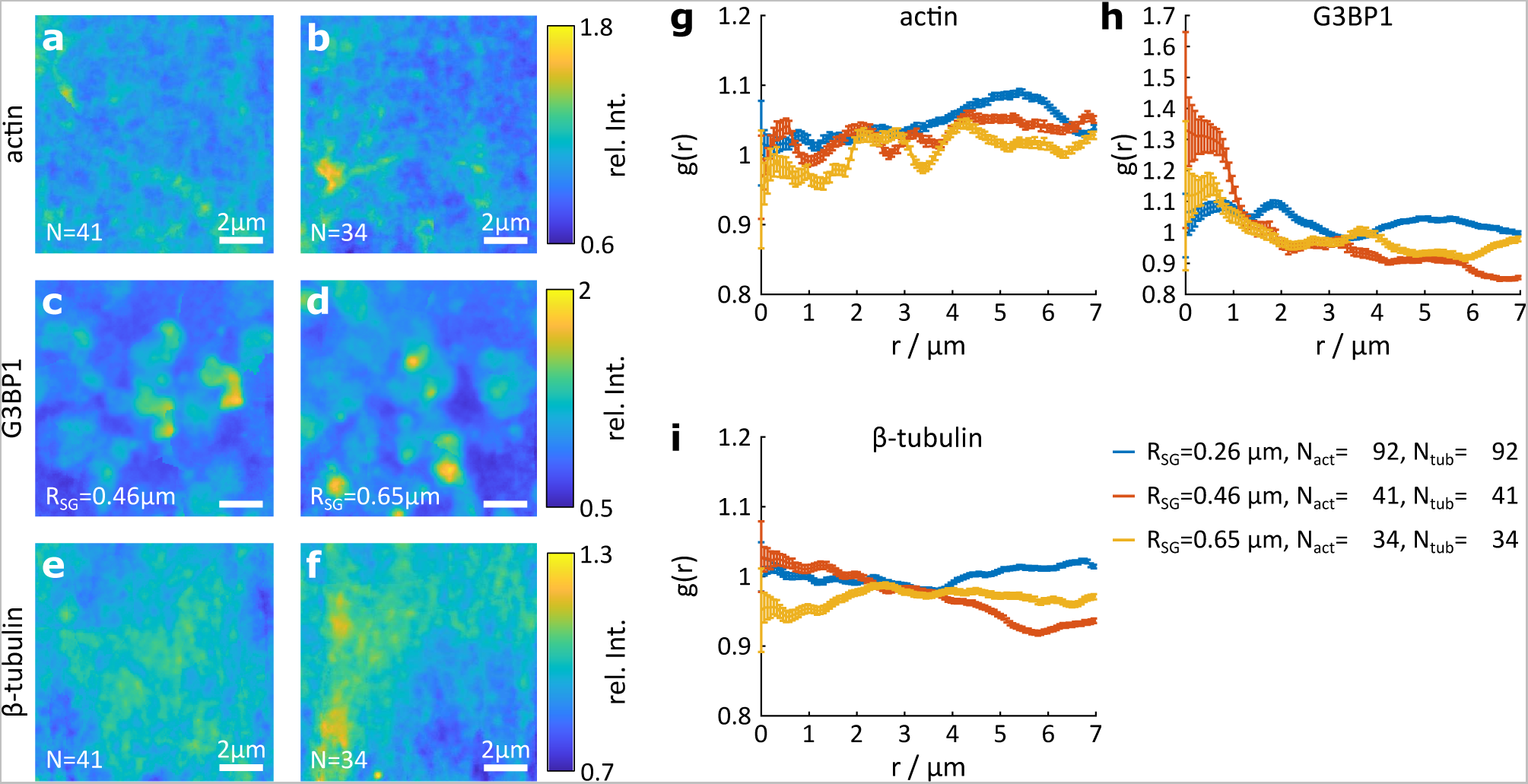
Negative control for distribution maps and *g*(*r*) after 30 min of blebbistatin treatment before 90 min of arsenite treatment, see Fig. 17. **a**, **b**, negative control for the distribution maps for actin for granules with *R_SG_* in the interval of [0.39 µm 0.52 µm] and [0.59 µm 0.72 µm] respectively. *N* gives the number of contributing SGs. **c**, **d**, the corresponding controls for G3BP1. The number of contributing SG is identical to those for actin. **e**, **f**, control for *β*−tubulin around SGs of the same size. Note that the number of contributing SGs is slightly different. **g**, *g*(*r*) of the control for actin around SGs of varying *R_SG_*. **h**, *g*(*r*) of the control for G3BP1 around SGs of varying *R_SG_*. **i**, *g*(*r*) of the control for *β*−tubulin around SGs of varying *R_SG_*.

## Footnotes

1 Note that SGs form a bit slower in U2OS wild-type cells with SGs forming typically within 10 to 20 min.

2 Using Fixed cell data and asking out the nucleus based on its 2D shape projected into 3D, we find an average volume of the cell cytosol of 1380 μm^3^, see Fig. 3 d.

